# Critical Negatively Charged Residues Are Important for the Activity of SARS-CoV-1 and SARS-CoV-2 Fusion Peptides

**DOI:** 10.1101/2021.11.03.467161

**Authors:** Alex L. Lai, Jack H. Freed

## Abstract

Coronaviruses are a major infectious disease threat, and include the human pathogens of zoonotic origin SARS-CoV (“SARS-1”), SARS-CoV-2 (“SARS-2”) and MERS-CoV (“MERS”). Entry of coronaviruses into host cells is mediated by the viral spike (S) protein. Previously, we identified that the domain immediately downstream of the S2’ cleavage site is the *bona fide* FP (amino acids 798-835) for SARS-1 using ESR spectroscopy technology. We also found that the SARS-1 FP induces membrane ordering in a Ca^2+^ dependent fashion. In this study, we want to know which residues are involved in this Ca^2+^ binding, to build a topological model and to understand the role of the Ca2+. We performed a systematic mutation study on the negatively charged residues on the SARS-1 FP. While all six negatively charged residues contributes to the membrane ordering activity of the FP to some extent, D812 is the most important residue. We provided a topological model of how the FP binds Ca^2+^ ions: both FP1 and FP2 bind one Ca^2+^ ion, and there are two binding sites in FP1 and three in FP2. We also found that the corresponding residue D830 in the SARS-2 FP plays a similar critical role. ITC experiments show that the binding energies between the FP and Ca^2+^ as well as between the FP and membranes also decreases for all mutants. The binding of Ca^2+^, the folding of FP and the ordering activity correlated very well across the mutants, suggesting that the function of the Ca^2+^ is to help to folding of FP in membranes to enhance its activity. Using a novel pseudotyped virus particle (PP)-liposome methodology, we monitored the membrane ordering induced by the FPs in the whole S proteins in its trimer form in real time. We found that the SARS-1 and SARS-2 PPs also induce membrane ordering as the separate FPs do, and the mutations of the negatively charged residues also greatly reduce the membrane ordering activity. However, the difference in kinetic between the PP and FP indicates a possible role of FP trimerization. This finding could lead to therapeutic solutions that either target the FP-calcium interaction or block the Ca^2+^ channel to combat the ongoing COVID-19 pandemic.

## INTRODUCTION

Since the outbreak of the ongoing COVID-19 pandemic in Wuhan, China at the end of 2019, there have been over 200 million cases and more than 4 million deaths associated with COVID. The United States alone has more than 35 million cases and 0.6 million casualties. It has severely harmed our public health, economy and social life. Even though vaccines have been developed, the rapid evolution of variants still pose a great threat: the delta variant can escape from the protection of most vaccines, and as a result, even fully vaccinated persons have a high possibility to be infected. A better understanding of the mechanism of COVID19 transmission is among the central agendas to save our society from the pandemic.

The cause of COVID-19 is SARS-2, which belongs to the family of coronaviruses, a diverse group of enveloped viruses that infect humans and animals, with several recent examples of zoonotic transmission including SARS-1 and MERS^1^, which were responsible for the pandemic outbreaks of) in 2003 (774 deaths) and 2012 (624 deaths), respectively^1^. These three viruses share a lot of common features, including the mechanism of viral infection.

The viral spike protein (S) is a CoV glycoprotein located on the viral envelope. It is required to mediate the viral entry into the host cells and is a major pathogenicity determinant^2, 3^. The S proteins of SARS-1, SARS-2 and MERS anchor on the viral envelop membrane in the form of trimers. Each monomer consists of S1 and S2 subunits. The two major steps in viral entry are 1) receptor binding, in which the S1 subunit recognizes a receptor on the host cell membrane, e.g., ACE2, and attaches the virus to the host cell, followed by 2) membrane fusion, in which the S2 subunit mediates the viral envelop membrane and the host membrane fusing together and releasing the virion into the host cell. Membrane fusion is a required stage in viral entry^4^; thus, the blocking of this membrane fusion could be an objective leading to vaccines and therapies to combat COVID-19^5^.

The major region of S that interacts with lipid bilayers of the host is called the “fusion peptide” (FP). Almost every enveloped virus has its glycoprotein and the corresponding FP, as it is critical for membrane fusion as it inserts into the host lipid bilayer upon activation of the fusion process, perturbing the membrane structure, and initiating membrane fusion^6, 7^. Previously, we used phospholipid (PC) spin labels to detect the local perturbation of the membrane by viral FP and TMD^8–14^.

The S proteins and FP of SARS-1, SARS-2 and MERS have several distinct features comparing to the other class I viral fusion protein, which includes the influenza hemagglutinin (HA) and the HIV envelope protein (Env). First, influenza and HIV envelope proteins are known to be activated via cleavage by host cell proteases to cleave the proto-glycoproteins into two subunits; this occurs at a single, restricted site directly upstream of their FP. In contrast, coronaviruses typically have two distinct cleavage sites (S1/S2 and S2’) that can be activated by a much wider range of proteases. As a result, we have shown that the FP is only exposed after the cleavage event at the S2’ site, and using ESR (Fig 1A) we have identified that the *bona fide* FP is directly downstream of the S2’ site for all of SARS-1, SARS-2 and MERS ^12, 15, 16^. This gives coronaviruses unique flexibility in the ability to invade new cell types, tissues and host species.

**Fig. 1.**
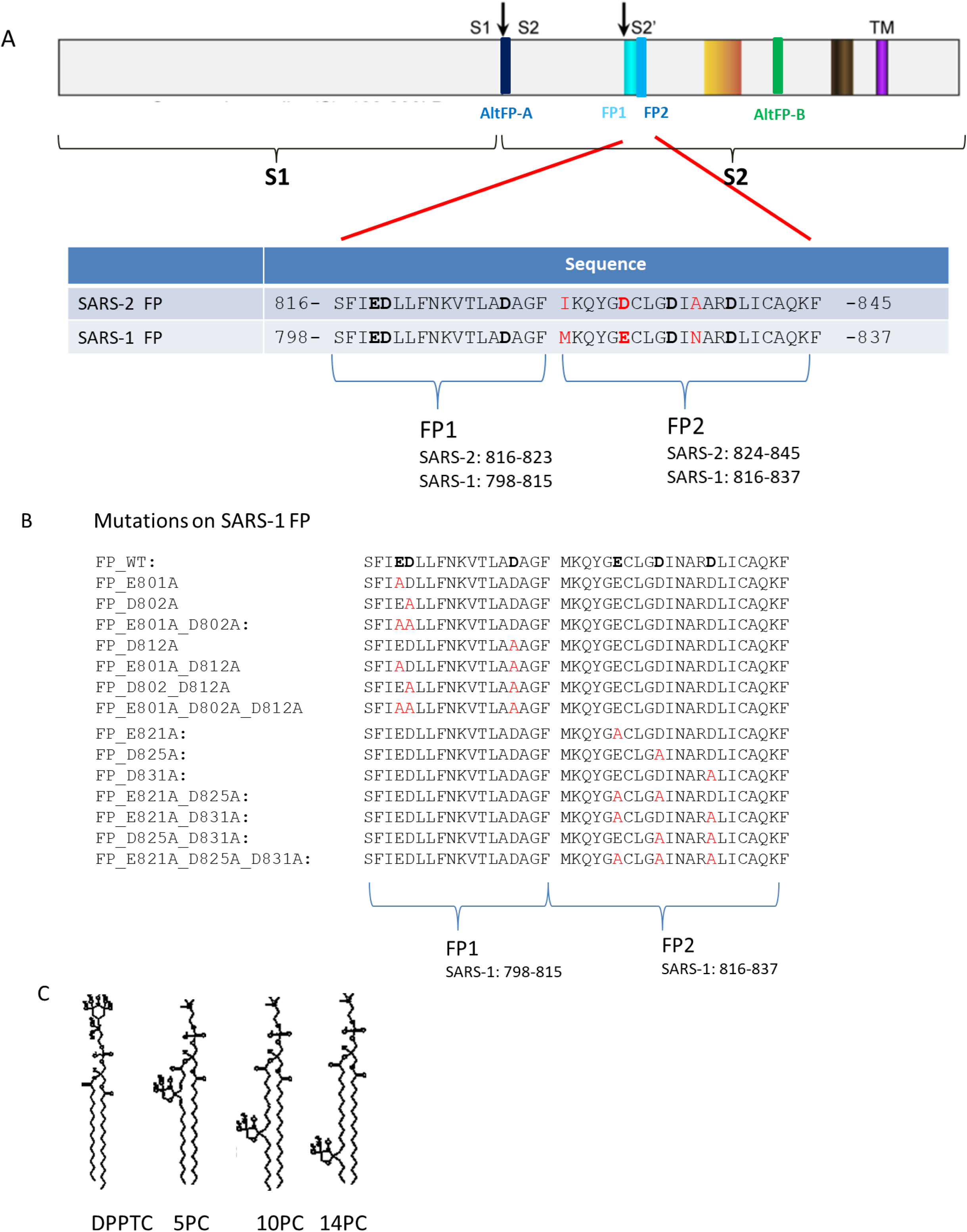
A) Diagram of subunits and domains of the S proteins of the SARS-1 and SARS-2 (upper panel) and the sequence alignment of the SARS fusion peptide domain (lower). The S protein is cleaved at the S1/S2 site into S1 and S2 subunits. The cleavage at the S2’ site exposes the N- terminal FP domain in the S2 subunit. We designate the FP1 and FP2 segment in the FP (previously designated as FP1_2) as shown in the lower panel. The residues in bold are the negatively charged residues. The residues in red highlight the difference between SARS-2 and SARS-1 FPs. B) The mutations for the SARS-1 FP used in this study. The residues in bold in sequence of the wild type (WT) are the negatively charged residues. The residue(s) in red indicates where the substitution occurs. C) The structure of spin labeled lipids used in this study: DPPTC, 5PC, 10PC and 14PC.

Second and more directly relevant to this study, we discovered the viral entry of the CoVs are Ca^2+^ dependent^12, 15, 16^, which is quite rare among the enveloped viruses^13^. We also discovered that the function of the CoV FPs is Ca^2+^ dependent, and while the SARS-1 and SARS-2 FPs bind Ca^2+^ at an 1:2 (FP:Ca^2+^) ratio, the MERS FP binds Ca^2+^ at an 1:1 ratio. Presumably, the FP is where the target for the Ca^2+^ in this Ca^2+^-dependent viral entry mechanism. This is important because the FP domain is very conserved comparing to other parts of the S protein, thus making it an ideal target for vaccine and antiviral drugs. It also raises the possibility of repurposing FDA-approved drugs in blocking Ca^2+^ channels for COVID treatment. However, how the CoV FPs interact with Ca^2+^ and how this interaction affects the function of the FP and the viral entry both remain unknown.

There is no crystal structure of intact S, severely limiting our mechanistic understanding of coronavirus fusion. While the basic structure of the S of SARS and SARS-2 can be revealed using cryo-EM techniques^17–20^ and part of the SARS-2 S2 subunit has been solved by X-ray crystallography^21, 22^, many structural and functional aspects remain undetermined. Furthermore, the FP adopts its active conformation only in the membranes^12^, which is generally not determined in the crystallographic structure as they are hydrophobic and intrinsically disordered. The structure of the SARS-2 FP has been solved only recently, using NMR in bicelles^23^. ESR is a useful technique to study the effect of FPs on the membrane structure with implications for the mechanism leading to membrane fusion. It can also be used to determine the peptide structure in the membrane in the form of Pulse-Dipolar ESR^24, 25^ and Power Saturation ESR^26^.

In this study we focus on the SARS-1 FP-Ca^2+^ interaction and its effect on membranes, using mainly ESR. We try to answer four questions in this research. First, as there are six negatively charged residues on the FP, which are involved in the Ca^2+^ binding? Second, what is the topology model of Ca^2+^ binding? Third, how is the Ca^2+^ binding enhanced the FP-induced membrane ordering? Fourth, does the separate FP function the same way as the FP on the whole S protein on the viral membrane does? For this purpose, we systematically introduced mutations on those residues, and examined their effect on membrane structure as well as their Ca^2+^ binding. We also used a more advanced pseudotype viral particle-SUV system to investigate the function of FP in the context of the whole protein, which better simulates the “biological scenario” ^13^ and compare the results with the those of separate peptides. In addition, we also show the study on a few corresponding mutants of the SARS-2 FP and prove this research on SARS-1 is helpful for the SARS-2 FP case.

## RESULTS

### Design of peptides and the lipid system

Viral FPs typically have low sequence conservation among different viral groupings. However, they are very well conserved within a given virus family. We previously demonstrated that the SARS-CoV S region immediately downstream of the S2’ cleavage site 798-835 (SFIEDLLFNKVTLADAGFMKQYGECLGDINARDLICAQKF) is the *bona fide* FP. We previously showed that the SARS-1 FP consists of FP1 and FP2 segments; the activity of SARS- 1 FP is Ca^2+^ dependent, with its FP1 and FP2 each binding one Ca^2+^ ion.^12^ (The FP2 segment is sometimes referred as FPPR^20^ in the case of the SARS-2 FP). Initially, we also designated the region 798-818 as FP1 and the region 816-835 as FP2 for the SARS-1 FP.^12^ However, the boundary between the FP1 and FP2 was arbitrary as the middle of the FP is a loose linker^27^; and in fact there is little overlap in our earlier designation of FP1 and FP2. Later, from homology comparison and experiments, we identified the corresponding region 816-845 as the FP for SARS-2^16^. In that paper, we designated the region 798-817 as FP1 and region 818-835 as FP2 for the SARS-1 FP, and the corresponding region 816-823 as FP1 and 824-845 as FP2 for the SARS-2 FP. We follow this convention in the paper (Figure 1A). There were six negatively charged residues in the SARS FPs, three in the FP1 region and three in the FP2 region. Those residues are potentially capable of interacting with the Ca^2+^. However, it is unclear whether they are really involved in Ca^2+^ binding. To examine this, we systematically synthesized the mutants with alanine substitution of those residues, and we made single, double, and triple substitutions in both the FP1 and FP2 regions as shown in Figure 1B.

The ESR signal of the spin labels attached to the lipids in membrane bilayers is sensitive to the local environment. Four spin labels were used: DPPTC has a tempo-choline headgroup and the spin is sensitive to changes of environment at the headgroup region; 5PC, 10PC and 14PC have a doxyl group in the C5, C10, or C14 position, respectively, of the acyl chain (Fig. 1C); and they are sensitive to the changes of local environment in the hydrophobic acyl chain region at the different depths. Using the NLSL software based on the MOMD model ^28, 29^, the order parameter of the spin can be extracted, which is a direct measure of the local ordering of the membrane. Thus, the effect of peptide binding on the structure of the membrane can be monitored. These four spin-labeled lipids have been used in previous studies, and their ability to detect changes in membrane structure has been validated ^30, 31^. Our previous studies examined the effect of various viral FPs, including those of influenza^8, 11^, HIV ^10^, Dengue Virus ^14^, Ebolavirus(EBOV)^13^, SARS- 1^12^, MERS^15^ and SARS-2^16^, as well as the FP of the ancestral eukaryotic gamete fusion protein HAP2 ^14^. All of these peptides were found to induce membrane ordering in the headgroup region as well as in the shallow hydrophobic region of bilayers (i.e., 5PC).

We used POPC/POPS/Chol=3/1/1 as the model membrane system to be consistent with previous studies^12^. PS is an anionic lipid that is found in the inner leaflet of the cytosolic membrane, and to some extent in endosomal membranes ^32^. It is critical for synaptic membrane fusion ^33–35^, and it also promotes viral entry of various viruses ^36, 37^. The PS was shown to be critical for HIV entry as well ^38^. This composition has been widely used in FP research ^39, 40^. It is also consistent with the system that we have used previously ^41^. We previously used MLV ^10, 11, 14^. We used SUV in this ESR study, which is better compared to our PP study described later in this work. And we found that the FP induced membrane ordering effect for these two systems are similar.

### The D812A in the FP1 segment has most reduced membrane ordering effect

We first studied the mutants in the FP1 segment. As shown in fig. 2A, when the peptide:lipid ratio (P/L ratio) of FP WT increases from 0 to 2 mol%, the order parameter S_0_ of DPPTC increases significantly from 0.44 to 0.48 at pH 5. The increase of the S_0_ is similar to the effect of influenza FP as we previously shown ^11^. All the three single mutants E801A, D802A and D812A have a lower membrane ordering activity than the WT. While the E801A only has a small reduced activity, the activity of D802A decreases a little more significantly. However, the activity of D812A drops the most of those three. In fact, its activity is very similar to that of the E801A_D802A double mutant, and only slightly higher than that of the E801A_D802A_D812A triple mutant. It is worthwhile to point out that even the triple mutant has a small ordering effect, which is due to two factors. First, the SARS-1 and SARS-2 has a “basal” activity even in the absence of Ca^2+^ as we have shown previously^16^. Second, even when all negatively charged residues in the FP1 segment are substituted, the FP2 segment remains unchanged; and we have shown previously that the FP2 alone also mediates membrane ordering, though in a smaller extent than the FP1^12^.

**Fig. 2.**
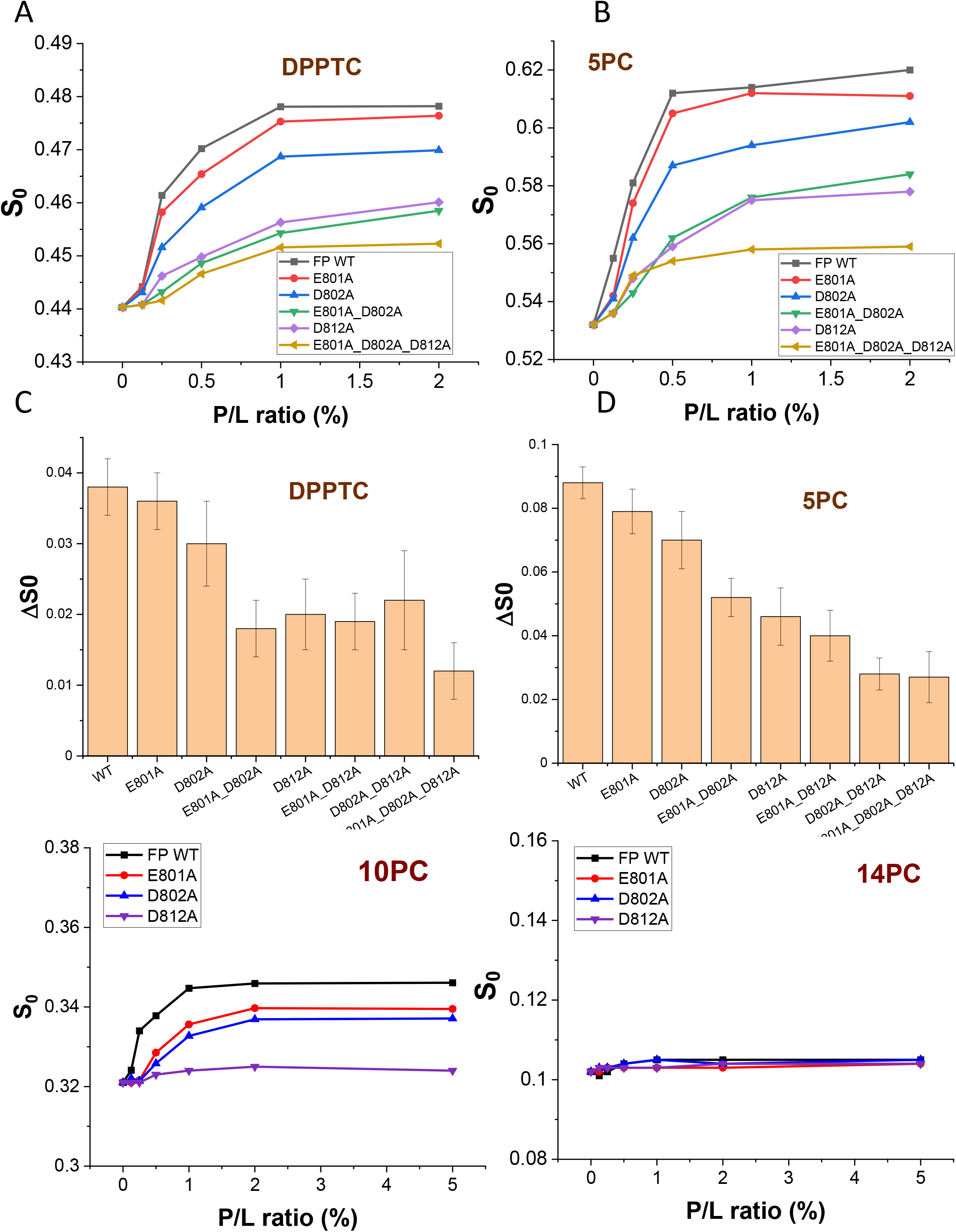
Membrane ordering for SARS-FP mutants in FP1 segment. A-B, E-F) Plots of order parameters of DPPTC (A), 5PC (B), 10PC (E) and 14PC (F) versus peptide:lipid ratio (P/L ratio) of SARS-1 FP in POPC/POPS/Chol=3/1/1 SUV in buffer 5 buffer with 150 mM NaCl at 25 °C. Black, WT; red, E801A; blue, D802A; green, E801A_D802A; purple, D812A, and yellow, triple mutants. C-D) The plot of ΔS_0_ for the WT and mutants of DPPTC (C) and 5PC (D) at P/L ratio=2%. The ΔS_0_ is calculated as S_0_(2% P/L)-S_0_(0% P/L). The experiments are typically repeated three times. The typical uncertainties we find for S_0_ range from 1-5 × 10^-3^, while the uncertainties from repeated experiments are 5-8 × 10^-3^ or less than ±0.01.

The membrane ordering of 5PC induced by the FP WT and its mutants also have the similar pattern (Fig 2B). The S_0_ significantly increases from 0.51 to 0.62 at pH 5 when the FP WT concentration increases. While all mutants have a reduced ordering activity, the D812A mutant exhibits the least active of those three single mutants. In fact, its maximal S_0_ at 2% P/L ratio is even slightly lower than that of the E801A_D802A double mutants, although the triple mutant has the lowest activity. Again, even the triple mutant has some “basal” activity as explained above. The reduction for 5PC is significantly greater than that for DPPTC. This is consistent with the fact that the ordering effect of the WT itself on 5PC is stronger than that on DPPTC. We suggest that the majority portion of the FP docks in the shallower hydrophobic region, and thus the membrane order in this region is the most sensitive to the FP. As a result, the effect of the mutation is also most significant in the 5PC region.

From the data of Fig 2A) and 2B), we plot the maximal ΔS_0_ which is calculated as ΔS_0_=S_0_ (at 2% P/L ratio)-S_0_ (at 0% P/L ratio) for both DPPTC and 5PC cases (Fig 2C and 2D, respectively). We also include the results of the other two double mutants E801A_D812A and D802A_D812A which have not been shown in Figs 2A and 2B. Those two double mutants have a membrane ordering activity similar to the D812A mutant in the headgroup region (DPPTC, Fig 3C), but their perturbation of the shallow hydrophobic region (5PC) is significantly smaller than that of D812A, especially for the D802A_D812A (Fig 2D).

**Fig. 3.**
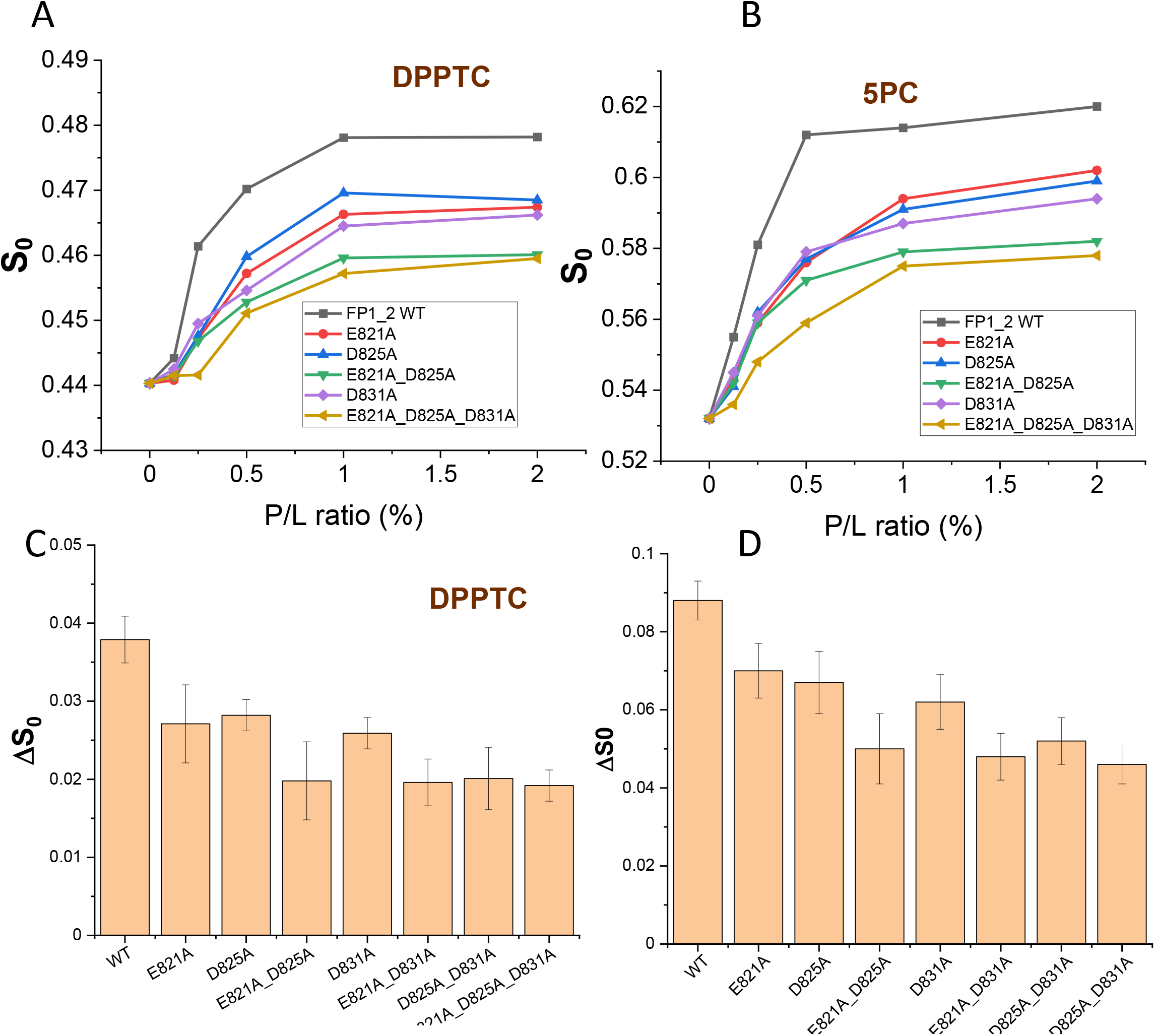
Membrane ordering for SARS-FP mutants in FP2 segment. A-B) Plots of order parameters of DPPTC (A) and 5PC (B) versus peptide:lipid ratio (P/L ratio) of SARS-1 FP in POPC/POPS/Chol=3/1/1 SUV in buffer 5 buffer with 150 mM NaCl at 25 °C. Black, WT; red, E821A; blue, D825A; green, E821A_D825A; purple, D831A; and yellow, triple mutants. C-D) The plot of ΔS_0_ for the WT and mutants of DPPTC (C) and 5PC (D) at P/L ratio=2%. The ΔS_0_ is calculated as S_0_(2% P/L)-S_0_(0% P/L). The experiments are typically repeated three times. The typical uncertainties we find for S_0_ range from 1-5 × 10^-3^, while the uncertainties from repeated experiments are 5-8 × 10^-3^ or less than ±0.01.

We also performed this experiment for the 10PC and 14PC, which corresponds to the middle hydrophobic region and deep hydrophobic region, respectively. The FP WT can induce membrane ordering at the 10PC region, while neither FP1 and FP2 can^12^. Thus, the membrane ordering effect in the middle hydrophobic region can be regarded as a cooperative effect between these two segments. As shown in Fig 2E, even though E801A has only slightly less ordering activity than that of the WT in the DPPTC and 5PC regions, its ordering activity reduces more significantly in the 10PC region. The D812A has basically no ordering effect to 10PC at all. The interaction of FPs with lipid bilayers has virtually no effect on the S_0_ of 14PC even for the WT (fig. 2F).

Thus, although all single mutants show a reduced ordering activity, the extents of the reduction are different. The mutated versions of the first two negatively charged residues, E801 and D802, have the smallest effect (Fig 2 A-D). Considering that they are next to each other’s locations, it is reasonable to suggest that their Ca^2+^ binding functions are redundant. When one has something else substituted for it, the other can assume most of its Ca^2+^ binding function. On the other hand, D812 is the most important residue. Its mutation causes the most substantial reduction of ordering activity of all single mutants. Its activity in the DPPTC and 5PC region is significantly lower than those of E801A and D802A, and in fact, is very close to the double mutant E801_D802 (Fig 2 A-D). The ordering activity of D812A is extraordinarily low in the region of 10PC (Fig 2E).

### All three negatively charged residues in the FP2 segment contribute to the membrane ordering effect equally

We then studied the mutants in the FP2 segment. As shown in fig. 3A, all the three single mutants E821A, D825A and D881A have a significantly lower membrane ordering activity than that of the WT. While the membrane ordering between them may show slight differences, they are at very similar levels, which is greater than that of E801A and D802A, but smaller than that of D812A. This is more clearly shown in the maximal ΔS_0_ plot (Fig 3C, again; ΔS_0_ is calculated as ΔS_0_=S_0_ (at 2% P/L ratio)-S_0_ (at 0% P/L ratio)). Interestingly, all three double mutants have a low ordering, nearly as low as the triple mutant. This pattern is different from the mutants in the FP1 segments, in which the triple mutant has a significant lower activity than the double mutants. This pattern is also reflected in the 5PC (Fig 3B and 3D). The three single mutants have a similar level of membrane ordering activity, which is lower than that of the WT. The double mutants and the triple mutants have a similar and even lower level of ordering activity. This strongly suggests that E821, D825 and D831 bind to an ion together and share the contribution almost equally. On the other hand, the ΔS0 value for the mutants in the FP1 regions (Figure 2C and 2D) does not have such a clear cut effect at 2% P/L ratio, suggesting the contributions of those three residues are uneven. All mutants basically have no ordering activity in the middle and deep hydrophobic regions (not shown).

Thus, we conclude that all three negatively charged residues in the FP2 segments contribute to the membrane ordering equally. This is different from the negatively charged residues in the FP1 segments. The D812 residue is obviously more important than either of the E801 or D802. In fact, its contribution to the membrane ordering activity is almost equal to the E801 and D802 combined, as its activity is similar to the E801A_D802A double mutant.

### Increased Ca^2+^ concentration promote the formation of secondary structures and the membrane ordering effect

To investigate why the membrane ordering effect of SARS-1 FP is Ca^2+^ dependent, we used circular dichroism (CD) spectroscopy to examine the secondary structure of the FP in membranes with different Ca^2+^ concentrations, which allows us to monitor the structural transitions occurring. As shown in fig. 4A, the FP has a greater ellipticity in the 1mM Ca^2+^ condition than in the 1mM EGTA condition at pH5 with SUV in 1:100 P/L ratio, indicating it contains more helical component. Using the H2D3 software^42^, we can estimate the percentage of alpha helix and beta strand components in the CD spectra. In the presence of 1mM Ca^2+^, it has 15.6% helix and 14.8% beta strand, and in the absence of Ca^2+^, it has 9.4% helix and 17.2% beta strand. We collected the CD spectra in various Ca^2+^ concentration, calculated the helical %, and plotted the helical % versus Ca^2+^ concentration (inlet of Fig 4A). We found that the helical concentration increases from 9% at 0 mM Ca^2+^ to 18% at 2% Ca^2+^, which is mostly saturated; and the X_50_ point, the concentration of Ca^2+^ to allow half of the maximal membrane ordering, is around 0.8 mM. Although the change is not dramatic, it is significant and indeed shows a clear trend. Thus, the Ca^2+^ promotes the folding of the FP in the membrane. This is consistent with the results we obtained from SARS-2 FP.^16^

**Fig. 4.**
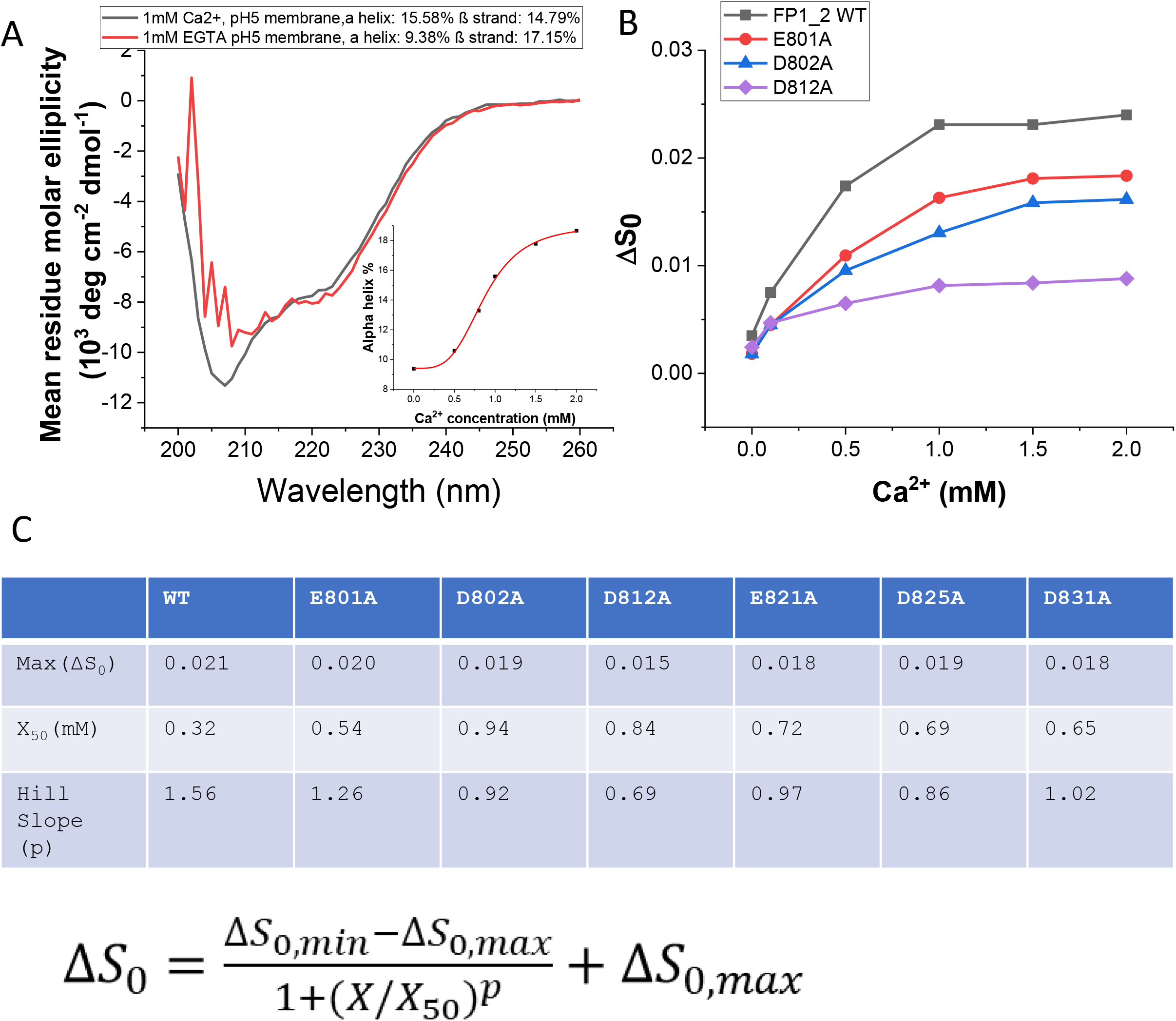
The effect of Ca^2+^ concentration. A) CD spectra of WT FP in membrane at 25 °C at pH5 with 1mM Ca^2+^ (black) and 1 mM EGTA (red). The inlet shows the alpha helix content % v Ca^2+^ concentration. B) The plot of different of order parameters of DPPTC between with and without peptide binding (ΔS_0_) versus Ca^2+^ concentration in POPC/POPS/Chol=3/1/1 SUV in pH5 buffer with 150 mM NaCl at 25 °C. Black, WT; red; E801A; blue 802A; and purple, D812A. C) The parameters of Max(ΔS_0_), X_50_, and Hill slope obtained from the fittings of (B) using the logistic function 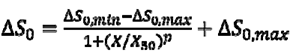.

### The mutations reduce the response to the Ca^2+^ in the ordering activity

To investigate the effect of the Ca^2+^ concentration, we repeated the ESR experiment using DPPTC. This time, however, we fixed the P/L ratio, increased the Ca^2+^ concentration from 0 to 2 mM, and extracted the S_0_. The highest calcium concentration used here is higher than the extracellular concentration of Ca^2+^ in human adult lungs (about 1.3 mM) ^43^, and from the CD experiments we know that 2mM Ca^2+^ has already saturated the structural change of the FP. The increase in S_0_ has two sources, the FPs and the Ca^2+^; thus, we generated a ΔS_0_-Ca^2+^ concentration plot, where ΔS_0_ = S_0_ (membrane with 1% FP) − S_0_ (membrane without FP) at each Ca^2+^ concentration. This subtraction cancels the membrane ordering induced by Ca^2+^ only (although it is relatively small), with the ΔS_0_ at each Ca^2+^ concentration representing only the contributions of the FPs. As shown in fig. 4B, Ca^2+^ increases the ΔS_0_ of the FP WT; we call it the “Ca^2+^ response effect”. This Ca^2+^ response effect gets saturated when the Ca^2+^ concentration exceed 1mM. This Ca^2+^ response effect reflects how the binding of Ca^2+^ of the FP promotes the folding of the FP in membrane and thus enables the FP to induce a greater ordering effect. We performed the same experiments on the six single mutants in both the FP1 and FP2 segments. If a mutant has a flatter Ca^2+^ response, then the corresponding mutated residue has a greater contribution in the Ca^2+^ binding. As shown in Fig 4B for the three mutants in the FP1 segment, D812A again shows a flat Ca^2+^ response compared to the WT; and the Ca^2+^ responses of E801A and D802A are flatter than that of the WT but still significantly steeper than that of the D812A. We fitted the curves with the logistic function 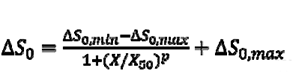 with fixed S_0,min_ and flexible S_0,max_, and also extracted the parameters of Max(ΔS_0_), X_50_ and Hill slope (p) from the curves (Fig 4C). The Max(ΔS_0_) indicates the maximal boost of the Ca^2+^ to the membrane ordering effect induced by to the FPs. As shown in Fig 4C, the Max(ΔS_0_) of D812A is only 0.015, which is much smaller than the WT (0.021). The Max(ΔS_0_) for the three single mutants in the FP2 segments are again very similar, which is consistent with the results showing that they contribute equally to the Ca^2+^ binding.

### The A811D substitution rescues the reduced activity of D812A

From the results above, we find that the D812 is a very important residue. We want to further understand whether this importance is related solely to its charge. Thus, we mutated a residue next to D812, A811 to D in both the WT (so we have the A811D) and the D812A (so we have the A811D_D812A) and repeated the ESR experiments (Figure 5A). The hypothesis is that if the charge is the only reason for its importance in Ca^2+^-dependent membrane ordering, then for the A811D_D812A the substitution of Asp for Ala at position 811 could “rescue,” at least partially, the “loss of Ca^2+^ binding function” of the substitution of Ala for Asp at position 812. And for th A811D, the two changed residues side by side will increase the binding of Ca^2+^.As shown in figure 5B and 5C, the ΔS_0_ of DPPTC significantly increases to 0.034 induced by 2% P/L ratio of the double mutant A811D_D812A, which is close to that induced by the same amount of WT (0.038). The A811D mutant actually induces an even greater ordering (ΔS_0_=0.040) than the WT, but the difference is not statistically significant. For the ΔS_0_ of 5PC, the double mutant rescues even more ordering activity than it does in the DPPTC case, as the ΔS_0_ is almost equal to that of the WT. The A811D mutant shows a little greater ordering activity than the WT in the case of 5PC, though the difference is not statistically significant either. We also repeated the ESR experiment for the Ca^2+^ response. As shown in Fig 5D, both the double mutant and A811D have a Ca^2+^ response similar to that of the WT, and both much greater than that of the D812A.Thus, the data suggest that the “loss of function” of the D812->A substitution can be largely rescued by A811->D substitution. The reason why the A811->D substitution cannot fully rescue the activity may be due to the unfavorable spatial position when D811 binds the Ca^2+^ ion compared to when the native D812 does. The fact that the A->D substitution (A811D) does not significantly increase the activity of the FP indicates that one charged residue is strong enough to bind to th Ca^2+^ from one location and the additional charged residue in the same location only has a marginal benefit.

**Fig. 5.**
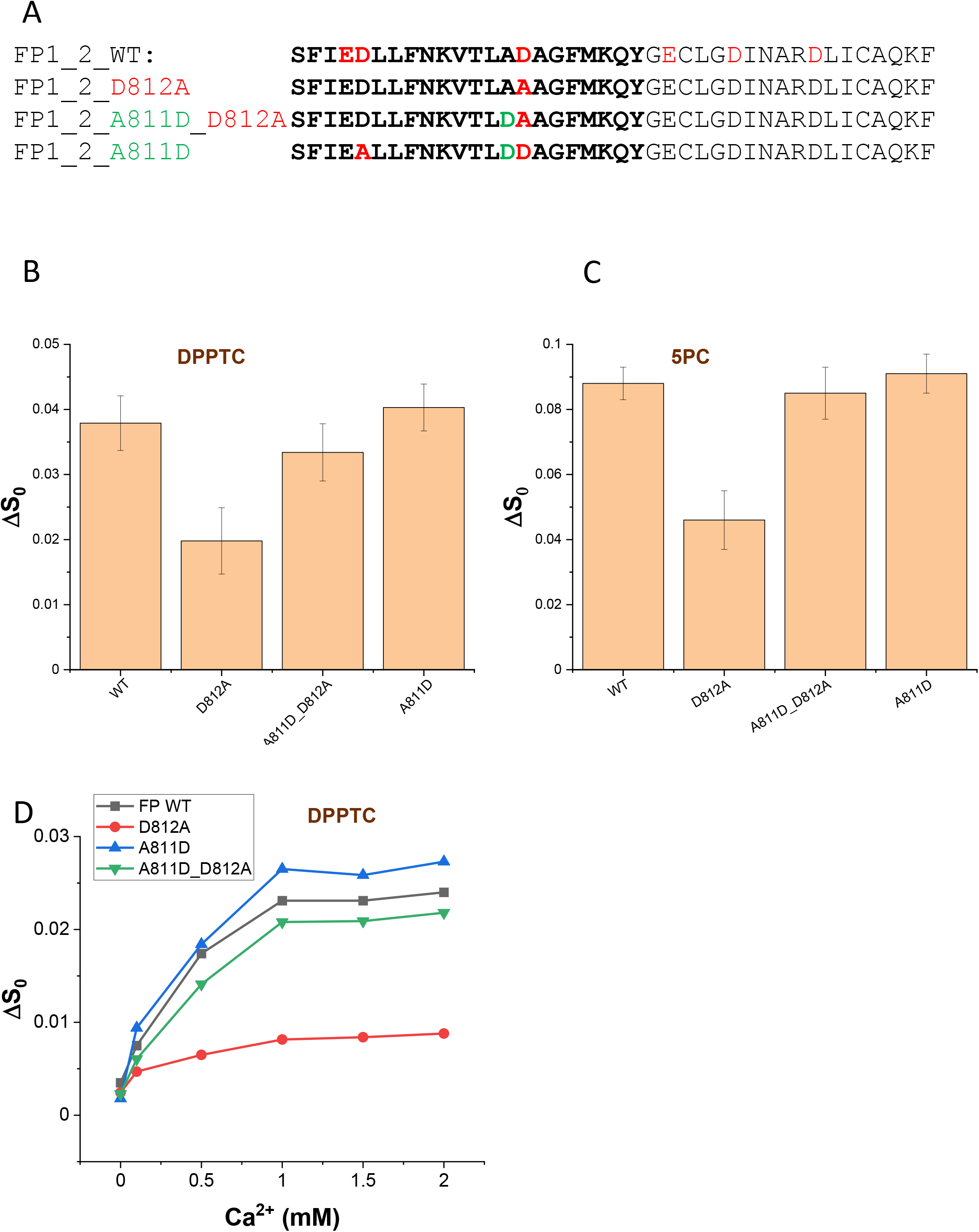
The substitution of A811D rescues the reduced activity of D812A. A) The sequence of WT FP, D812A, A811D and A811D_D812A. The residue in red highlights the D812A, and the residue in green highlights A811D. B-C) The plot of ΔS_0_ of DPPTC (B) and 5PC (C) for the WT and mutants of DPPTC. D) The plot of different of order parameters of DPPTC between with and without peptide binding (ΔS_0_) versus Ca^2+^ concentration in POPC/POPS/Chol=3/1/1 SUV in pH5 buffer with 150 mM NaCl at 25 °C. Black, WT; red; D812A; blue A811D; and green, A811D_D812A.

### Interactions of FPs with Calcium Cations Detected by ITC

We used ITC to investigate whether Ca^2+^ cations directly interact with SARS-CoV FPs. During titration, CaCl_2_ was injected into a reaction cell containing FPs. The background caused by th dilution of CaCl_2_ has been subtracted by using data from a control experiment that titrated CaCl_2_ in a pH 5 buffer. Substantial heat absorbed during the titration was observed, with the heat absorbed saturating toward the end of the titration. The data were fit using a one-site model, which hypothesizes that all binding sites have the same binding coefficient. While this the two binding sites may not have the same binding coefficient, especially for some mutants, this model can be used to estimate the “average” binding coefficient (K_b_) and N (Ca^2+^ to FP ratio).

As shown in fig. 6A, when the heat vs. molar ratio plot was fitted using a one-site model that assumes that all binding sites have the same binding affinity, we calculated the enthalpy change ΔH=6.77 kCal/mol, with binding constant K_b_=2.49×104 M^-1^ and stoichiometry N=1.70. From these parameters, we further calculated the free energy change ΔG=−RTlnK_b_=6.49 kCal/mol and −TΔS= ΔG−ΔH = −1.80 kCal/mol. The data show that the calcium-FP interaction is endothermic, and the binding ratio is close to two calciums per peptide. The reason for the endothermic reaction could be because the buffer solution (150 mM NaCl) contains significant amounts of Na^+^ ions that could also interact with FP and that a stronger binding of Ca^2+^ cations displaces bound Na^+^ into the bulk solution. The ITC experiments we conducted demonstrate strong evidence for direct calcium-FP interactions. We performed this experiment for the mutants as well. As shown in Fig 6B, the D812A mutant has a very similar ΔH as the WT, and the binding stoichiometry N=1.75 is not so different from that of the WT as well. However, it has an obviously flatter slope, which reflects a weaker binding constant. The triple mutant E801A_D802A_D812A also has an even flatter slope (Fig 6C), and its enthalpy is significantly smaller (5.97 kCal/mol); and most importantly, the binding stoichiometry N=1.19, which is very close to that of MERS FP. MERS FP has known to bind only one Ca^2+^ ion, not two^15^. Thus, the triple mutant only binds one Ca^2+^, which is consistent with the fact that all negatively charged residues in FP1 segment are substituted. We repeated the experiments for other mutants and list the parameters in Fig 6D. The triple mutant in the FP2 segment gives N=1.20, which is also close to that of the MERS FP (i.e., has only one binding site).

**Fig 6.**
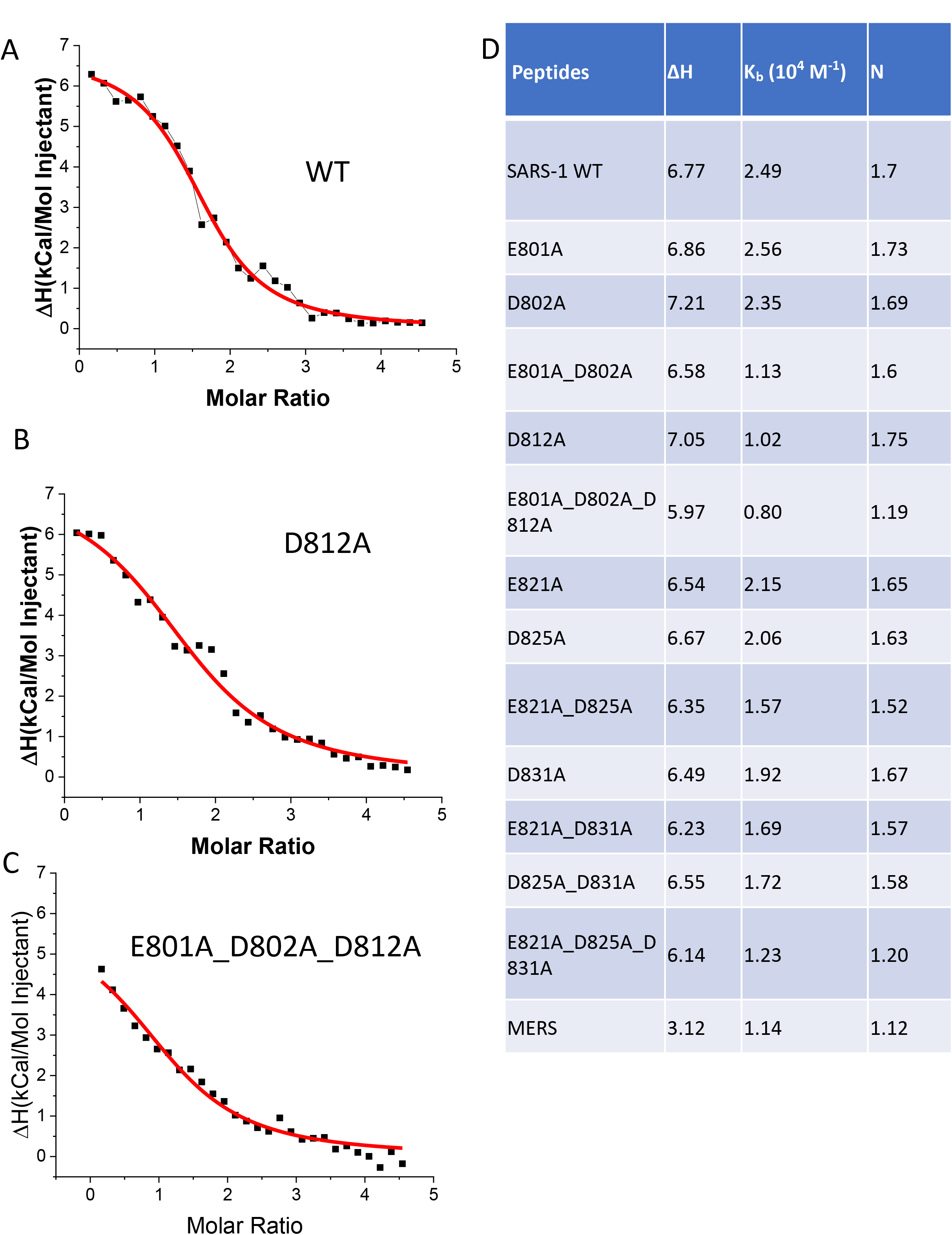
ITC analysis of Ca^2+^ binding to FP of the coronaviruses. (A) SARS-1 FP, (B) D812A, (C) triple mutant (E801A_D802A_D812A). The peptides were titrated with CaCl_2_ in the pH5 buffer. The integrated data represent the enthalpy change per mole of injectant, ΔH, in units of kcal/mol as a function of the molar ratio. The data were fitted using a one-site model. Data points and fitted data are overlaid. D) parameters of the FP-Ca^2+^ titration obtained from the ITC curves using one-site model.

### Enthalpy Changes of FP and Membranes Interaction

As we have shown in the CD experiment, the Ca^2+^ promotes the folding of the FP in the membranes. The folding of the FP can be further analyzed by comparing membrane binding enthalpies of FPs measured by ITC. To measure the binding enthalpy, small amounts of peptide were injected into a reaction cell containing a large excess of SUV lipid membranes. Thus, during the whole titration process the amount of available membrane can be regarded as constant, and all injected peptides can be regarded as binding to the membranes. As a result, the reaction heat in each injection is equal. The enthalpy of reaction can be calculated from the average of heat in each injection ^44, 45^. The FP-membrane participation is driven by enthalpy and opposed by the entropy as previously shown^46^. We performed the experiments under two conditions, in the presence of 1mM Ca^2+^ and in the presence of 1mM EGTA. In both cases, the same concentration of Ca^2+^ or EGTA are present in both the reaction cell and the injection syringe. Thus, there is no heat generated due to the Ca^2+^-peptide or the Ca^2+^-membrane interaction. Therefore, the heat detected is only generated from the membrane-FP interaction. The enthalpy gain (more negative enthalpy) is largely due to the formation of hydrogen bonds when the FP adopts a relatively fixed secondary structure when transferred from the solution (in which it is largely unstructured) to the membranes. A more negative enthalpy during the membrane binding reflects a better folding in the membrane.

We first examined the ΔH in the presence of Ca^2+^. As shown in Fig 7A, the WT folds the best in membranes as it has the greatest enthalpy gain (most negative), while the triple mutants E801A_D802A_D812A and E821A_D825A_D831A fold worst in membranes. If we compare all single mutants, the D812A again has significantly smaller enthalpy gains than either the E801A and D802A, which is consistent with the previous ESR results showing that this mutant has the lowest membrane ordering activity and lowest Ca^2+^ response. By comparing the ΔHs in the presence and absence of Ca^2+^, we found a pattern that for all WT and mutants, the enthalpy gain ΔH in the presence of Ca^2+^ is greater (more negative) than that in the absence of Ca^2+^. This indicates that the FPs fold better in membranes in the presence of Ca^2+^ in general. We also found that the ΔH and mutations are correlated well in both conditions, i.e., if a mutant has a greater enthalpy gain in the presence of Ca^2+^, it will have a greater enthalpy gain in the absence of Ca^2+^. We further calculated the difference of ΔHs between these two conditions and obtained the ΔΔH=ΔH(w/Ca)-ΔH(w/o Ca) as shown in Fig 7B. This ΔΔH represents how much better folding of the FP in the presence of Ca^2+^. The greater ΔΔH (more negative) indicates the greater effect of Ca^2+^ to promote the folding of the FP in membranes, which is related to the CD (Fig 4A) and Ca^2+^ response results investigated by ESR (fig 4B). We found that the two triple mutants have the smallest ΔΔH (−2.1 and -1.5 kCal/mol, respectively), while the WT has the greatest ΔΔH (−9.9 kCal/mol). This indicates that WT folds in the membrane much better in the presence of Ca^2+^ than in the absence of Ca^2+^, suggesting that the Ca^2+^ promotes the folding of WT most effectively. For all single mutants, D812A has the smallest ΔΔH (−5.1 kCal/mol), suggesting Ca^2+^ does not promote its folding as well as the other single mutants, which further indicates the importance of D812 in the FP-Ca^2+^ interaction.

**Fig 7.**
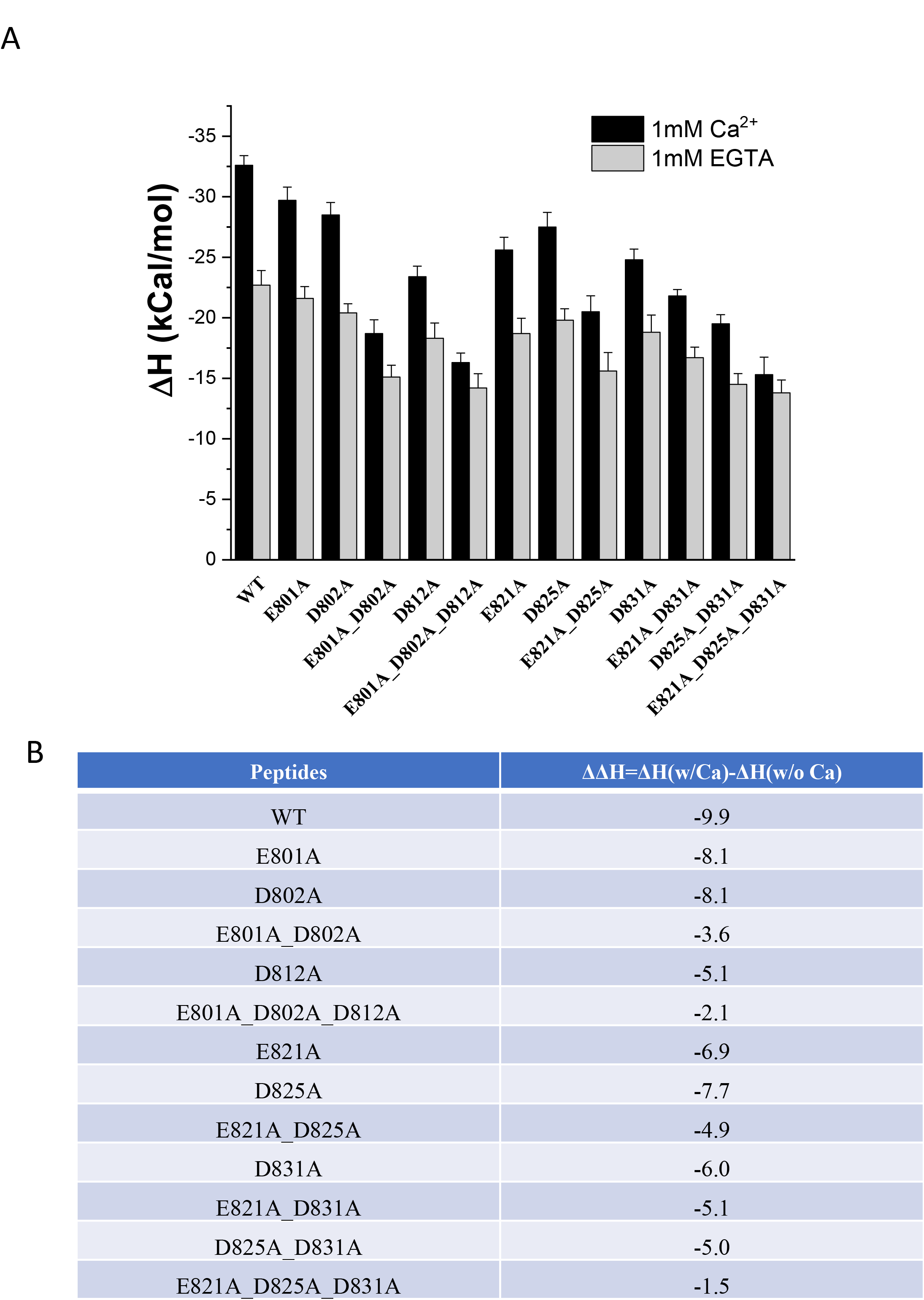
ITC analysis of FP-membrane interaction. A) enthalpy of SARS-1 FP-membrane interaction. The peptides were injected into the reaction cells containing POPC/POPS/Chol=3/1/1 SUV in buffer 5 buffer with 150 mM NaCl at 25 °C in the presence of either 1 mM Ca^2+^ (black) or 1mM EGTA (grey). The reaction heats of each injection were averaged and the reaction enthalpy was calculated. B) the ΔΔH of the WT and mutants, in which ΔΔH =ΔH(w/Ca^2+^)-ΔH(w/EGTA). This value indicates the difference of membrane binding enthalpy due to the presence of Ca^2+^.

### The mutants of SARS-2 FP show a trend similar to the corresponding SARS-1 FP mutants

We initially started this study long before the COVID19 outbreak. After the outbreak began, we immediately started to work on SARS-2 FP as well and we found that the SARS-2 FP also induces membrane ordering in a Ca^2+^ dependent fashion and its effect is even greater than that of the SARS-1 FP.^16^ While we focus on SARS-1 FP in this paper, we are performing parallel research on SARS-2 FP in more detail. The sequence of SARS-2 FP only has a three-amino-acid-difference from that of the SARS-1 FP (Fig 8A) and only one of them is negatively charged, which is E821 in the SARS-1 FP and the corresponding D839 in the SARS-2 FP (both of them are negatively charged). Thus, the knowledge we obtained from this research on SARS-1 FP is mostly applicable to SARS-2 FP as well. As shown in Fig 8B and Fig 8C, D839A (corresponding to E821A in the SARS-1 FP) has a smaller membrane ordering effect than the SARS-2 FP WT in both DPPTC and 5PC cases. And the D830A (corresponding to D812A in the SARS-1 FP) has an even smaller effect than that of D839A. These results are consistent with our findings in the SARS-1 FP on D812A and D821A.

**Fig. 8.**
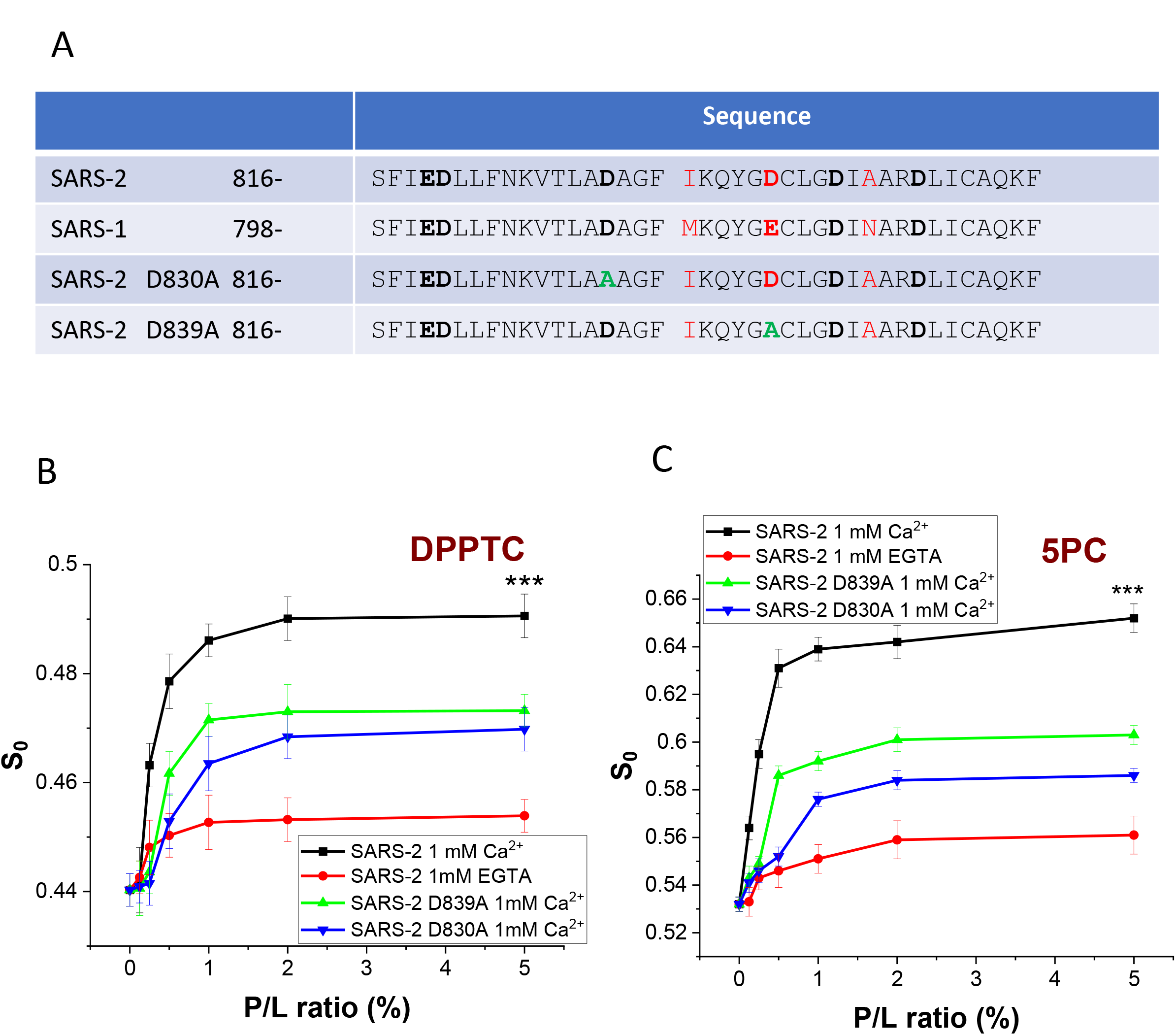
Membrane ordering of SARS-2 mutants. A) the sequence of SARS-2 mutants. B-C) Plots of order parameters of DPPTC (B) and 5PC (C) versus peptide:lipid ratio (P/L ratio) of SARS-2 FP in POPC/POPS/Chol=3/1/1 SUV in buffer 5 buffer with 150 mM NaCl at 25 °C. Black, WT in the presence of 1 mM Ca^2+^; green, D830A (corresponding to D812A in SARS-1 FP); blue, D839A (corresponding to E821A in SARS-1 FP); and red, WT in the presence of 1 mM EGTA.

### Ca^2+^ promotes insertion of viral S protein trimer into the target membrane in the PP-SUV docking system

It has been questioned whether the FP by themselves function the same as the FP in the viral glycoprotein. To address this issue and to better simulate the “biological Scenario” we have developed a methodology that we called “PP-SUV docking system” (Fig 9A) ^13^. The PPs are non-transmittable chimeric viral particles expressing viral glycoproteins such as SARS-1 S trimers on the surface of a replication-deficient murine virus. The PP with the S protein is pretreated by proteinase. When the PP is mixed with the SUV containing DPPTC, the SARS-1 FP will insert into the SUV membrane during the docking of the PP on the SUV. After mixing the PP and the SUV containing spin labeled lipids and triggering with Ca^2+^, we collected ESR spectra every 30 sec. In order to do that, we needed to reduce the scanning times which, however, yield noisier signals. These noisier signals were then denoised using the recently developed Wavelet Denoising Package^5^ before NLSL analysis, as required. We extracted the local S_0_ from each spectrum and plotted S_0_ vs. time^13^ (Fig. 6B-D). Using this methodology, we monitored in real time how the local membrane ordering develops in the SUV membranes as mediated by the FPs in the whole glycoproteins that are assembled on the PP surface: this better simulates the “biological scenario.”^13^

**Fig. 9.**
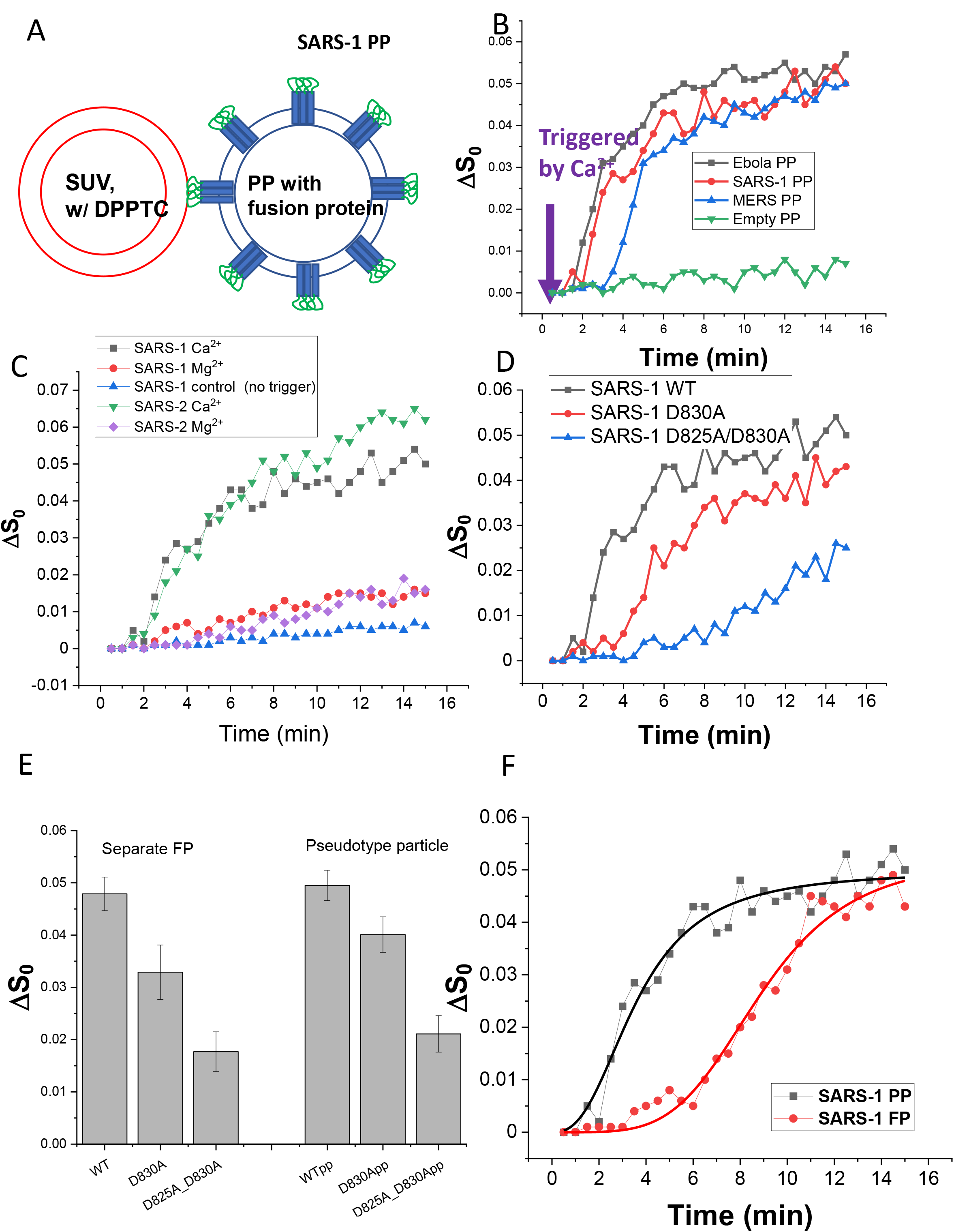
Time-dependent ESR experiments in the pseudotype viral particle (PP) - SUV docking system. A) the schematic diagram of the PP-SUV docking system. The fusion protein (in this case EBOV G protein or SARS-1 S protein) were expressed and assembled as trimers on the PP membrane. DPPTC was incorporated in POPC/POPS/Chol=3/1/1 SUV. The PP and SUV were mixed at a PP:SUV ratio of ca. 1:5, and triggered by the addition of 2 mM Ca^2+^. (B) the plot of local order parameters of DPPTC changes (ΔS_0_) during the time course of measurement. Black, EBOVpp; red, SARS-1 PP; blue, MERS PP; green, empty PP (no viral glycoprotein expressed). (C) Same as (B) using SARS-1 PP and SARS-2 PP in different triggering conditions. Black SARS-1 PP triggered by 2mM Ca^2+^; red, by 2mM Mg^2+^; blue, control by injecting the same volume of buffer; green, SARS-1 PP triggered by 2mM Ca^2+^; purple, by 2mM Mg^2+^. (D) Same as (B) using wildtype (WT) SARS-1 PP (black), D830A mutant (red), and D825A_D830A double mutant (blue). (E) comparing the ΔS_0_ induced by separated FP and PP. (F) comparing the time course of SARS-1 PP (black) and SARS-1 FP (red). In the FP case, we mixed the separate FP and the SUV: the amount of the SUV is the same as the one used in the PP case, and the amount of the FP is 1% of the amount of lipids. The PP has an early start and a steeper slope, while the FP has a late start and a flatter slope although they reach almost the same final ΔS_0_.

We used EBOVpp as a positive control as it was shown to work well^13^. We fitted the curve using a Boltzmann equation 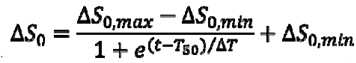, and obtained three parameters from each curve. The ΔS_0(max)_ is the maximal ordering occurs at the end of the time course, indicating the overall ordering activity. The T_50_ is the time to reach its 50% maximal ordering effect, indicating how early the PP starts to induce ordering. The ΔT is the slope at T_50_, indicating the rate of the change of ΔS_0_, which is how fast the PP induces the ordering once it starts to take effect. The parameters are summarized in Table 1.

**Table 1.** The parameters of fittings in pp-mediated membrane ordering. The curves are fitted using the Boltzmann function 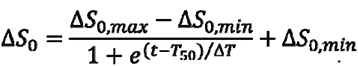

| | $\Delta S_{0,max}$ | $T_{50}$ (min) | $\Delta T$ (slope) (min<sup>-1</sup>) |
| --- | --- | --- | --- |
| EBOVpp | 0.051 | 3.2 | 1.02 |
| SARS-CoVpp | 0.049 | 3.6 | 1.19 |
| MERS-CoVpp | 0.045 | 4.8 | 0.92 |
| SARS-CoV-2pp | 0.058 | 3.8 | 1.51 |

B
| | $\Delta S_{0,max}$ | $T_{50}$ (min) | $\Delta T$ (slope) (min<sup>-1</sup>) |
| --- | --- | --- | --- |
| SARSpp WT | 0.049 | 3.6 | 1.19 |
| D830A | 0.040 | 5.6 | 1.61 |
| D825A_D830A | 0.035 | 12.3 | 2.98 |

C
| | $\Delta S_{0,max}$ | $T_{50}$ (min) | $\Delta T$ (slope) (min<sup>-1</sup>) |
| --- | --- | --- | --- |
| SARS-1 PP | 0.049 | 3.6 | 1.19 |
| SARS-1 FP | 0.051 | 8.9 | 4.47 |

As shown in Fig 9B with DPPTC, for the SARS-1 PP, ΔS_0_ increases in the presence of Ca^2+^, and the T_50_ occurs at about 3.6 min. If an empty PP (no viral glycoprotein on the surface) was used, there is no increase of S_0_. Thus, the increase in S_0_ observed is due to the S protein on the SARS-1 PP. The maximal ΔS_0_ of SARS-1 PP is the same as that obtained with just the separate FP (cf. Fig 2A and 2C and Fig 9E) and slightly smaller than that of the EBOV PP for which the T_50_ is slightly smaller (T_50_=3.2 min). The ΔT of SARS-1 PP is also slightly greater than EBOV PP (1.19 min^-1^ vs 1.02 min^-1^). However, this could result from the differences in protein expression level, surface transportation efficiency of the glycoprotein, and/or efficiency of proteinase cleavage. The MERS PP has a significantly slower start (T_50_=4.8 min), and once it starts to induce ordering, its ΔT is also slightly smaller (0.92 min^-1^).

We have previously shown that the Ca^2+^ dependence of SARS-2 FP is very specific^16^. We want to test whether this is the case for PP. We compared the triggering effect of Ca^2+^ and Mg^2+^. A shown in fig 9C, the injection of Mg^2+^ instead of Ca^2+^ triggers the ΔS_0_ slightly (ca. 0.015 at 15 min) while the Ca^2+^ triggers the ΔS_0_ up to ca. 0.049. There is a signature S-shape “jump” in the ΔS_0_-time curve in the Ca^2+^ case, whereas this shape is not observed in the Mg^2+^ case. If no trigger is added (the same volume of buffer is added instead), there is virtually no change in the ΔS_0_. These results show that Ca^2+^ is required for the whole S protein on the membrane fusion as expected. We previously noted that there is some “basal activity” for the isolated FP in the presence of EGTA^12, 16^ (also see Fig 8A), but the “basal activity” for the whole S protein seems to be negligible as shown in Fig 9C.

We recently obtained SARS-2 PP from Dr. Gary Whittaker Lab and Dr Susan Daniel Lab and performed a parallel PP study on it as the SARS-1 PP. As shown in Figure 9C and table 1A. The SARS-2 pp induces an even higher ordering effect than the SARS-1 PP, with a ΔS_0(max)_=0.058 comparing to SAR-1 PP’s 0.049. While it starts the “jump” a little more slowly than the SARS-1 PP, it has a faster rate (ΔT) once it starts to take effect. This is consistent with our previous research that the SARS-2 FP has a higher membrane ordering effect than the SARS-1 FP even though there are only three residue differences between them^16^. We also tested the Mg^2+^ as a trigger reagent. As expected, Mg^2+^ cannot trigger the SARS-2 PP to mediate membrane ordering as the Ca^2+^ does. The result confirm that that the Ca^2+^ dependence is very specific, as shown previously for the SARS-2 FP^16^ even for the PP.

We then studied SARS-1 PP mutations. As shown in fig 9D and Table 1B, the D830A single mutant does exhibit somewhat lower activity than the WT in two ways. First, its ΔS_0_(max) (0.040) is less than that of the WT (0.049). Second, its time development (T_50_=5.6 min) is slower than that of the WT (T_50_=3.6 min). The D825A_D830A double mutant has even lower activity. It has an even smaller ΔS_0(max)_ (0.035) at the end of our measurement and an even slower time development (T_50_=12.3 min). This result suggests that both D830 and D825 are involved in Ca^2+^ binding. The maximal ΔS_0_ of the separated WT and mutant FP (from experiments shown in figure 3A) and the WT and mutant PP (from Fig 9D) are compared side by side in Fig 9E. We can clearly see that the separate FP and the PP induce the ordering in a very similar level. The single mutant of both FP and PP induce a lower ordering, and the double mutant of both FP and PP induce even lower ordering. This result confirms that the knowledge that we obtained from the separate FP mutants can be largely translated to the FP in the context of whole protein in a trimer form anchored on the membrane.

However, we find that separate FP functions exactly the same way as the PP: the kinetic of how the FP and PP induce membrane ordering seems to be different. As shown in Fig 9F, we compare the FP induced membrane ordering in a time-dependent experiment with the PP. In this experiment, we mixed the FP with SUV in 1% P/L ratio, triggered by the injection of 2mM Ca^2+^ and collected the ESR spectra like the procedures that we used in the PP-SUV experiment. As we can see, although the ΔS_0(max)_ for the FP (0.051) are approximately the same as that for the PP (0.049), it took much longer to reach the maximal S_0_. The T_50_ estimated to be 3.6 min for the PP and 8.9 min for the FP (Table 1C). And even after the FP starts to perturb the membrane, the rate (ΔT=4.47 min^-1^) is much slower than that of the PP (ΔT=1.19 min^-1^). Although a careful calibration is required to reach a final conclusion, this likely suggests that the FPs in the whole protein trimer induce the membrane ordering more efficiently than the separate FP.

## DISCUSSION

### Which residues contribute to the Ca^2+^ binding?

The mechanism of membrane fusion is still only partially understood. Our past studies have shown that numerous viral FPs induce increased membrane ordering in a collective fashion, i.e., a significant increase in S_0_ as a function of the P/L ratio8,9,^11–15, 47^. We further suggested that FP-induced membrane ordering is accompanied with dehydration resulting from peptide insertion, and this is a prerequisite step for removal of the repulsive forces between two opposing membranes, thereby facilitating initialization of membrane fusion ^8–11, 14^. In conjunction with our collaborators, have also determined that the CoVs (SARS-1, SARS-2 and MERS) and EBOV require Ca^2+^ for viral entry, and that the Ca^2+^ binding site is on the FP^12, 13, 16^.

Our current study is the first to investigate the Ca^2+^ binding mechanism of CoV FPs in detail at the molecular level using biophysical methods. Although the negatively charged residues should be related to Ca^2+^ binding, the issues of which residues are involved and how much is its contribution remains unclear. In fact, in the case of MERS^15^ and EBOV^13^, not all negatively charged residues are important in the Ca^2+^ binding.

Our results suggest that all negatively charged residues are involved in Ca^2+^ binding, though their contributions are different. While those residues in FP2 (E821, D825 and D831) have equal contributions, the contribution of those in FP1 are unequal. E801 and D802 seems to share a redundant Ca^2+^ contribution. D812 is the most important single residue of all six. The extraordinarily low ordering activity of D812A in the middle hydrophobic region (10PC) is worth of more discussions. FPs such as that of HA are unable to induce membrane ordering as deep in this region. Neither FP1 and FP2 of SARS-1 can induce membrane ordering in this region as well^12^. On the other hand, we have observed a combination of peptides such as the HA FP-TMD dimer can induce membrane ordering in this region, suggesting the ability to induce ordering in this region represents a greater perturbation to the membranes and presumably promotes membrane fusion more efficiently^11^. Thus, the ability of the SARS-1 and SARS-2 FPs to induce membrane ordering at the 10PC region is special and may be important. Therefore, the lack of ordering effect in the 10PC region further indicates the importance of D812. The loss-of-function mutation D812A can be “rescue” by the mutation of a nearby A->D substitution at residue 811, indicating that the Ca^2+^ binding function of D812 is solely due to its charge.

### What is the Ca^2+^ binding topology?

Even though we have shown that both the separate FP1 and FP2 peptides bind one Ca^2+^^12^, this may not be the case when they are in the context of the full length FP (a.k.a. FP1_2). There could be at least several topological models with different topologies (Figure 10A). In first model, FP1 and FP2 each bind one Ca^2+^ by its own (Fig 10A left). Second, the negatively charged residues near the boundary between FP1 and FP2, namely D812 on FP1 or E821 on FP2 in SARS-1 FP, are involved in the binding of the “ion in the other side”, i.e., D812 binds an ion in the C-terminal (second Ca^2+^ ion), or E821 binds the ion in the N-terminal (first Ca^2+^ ion) (Fig 10A middle showing the former case). Third, the negatively charged residues in the middle bind one ion, and those in both termini bind the other (Fig 10A right).

**Fig 10.**
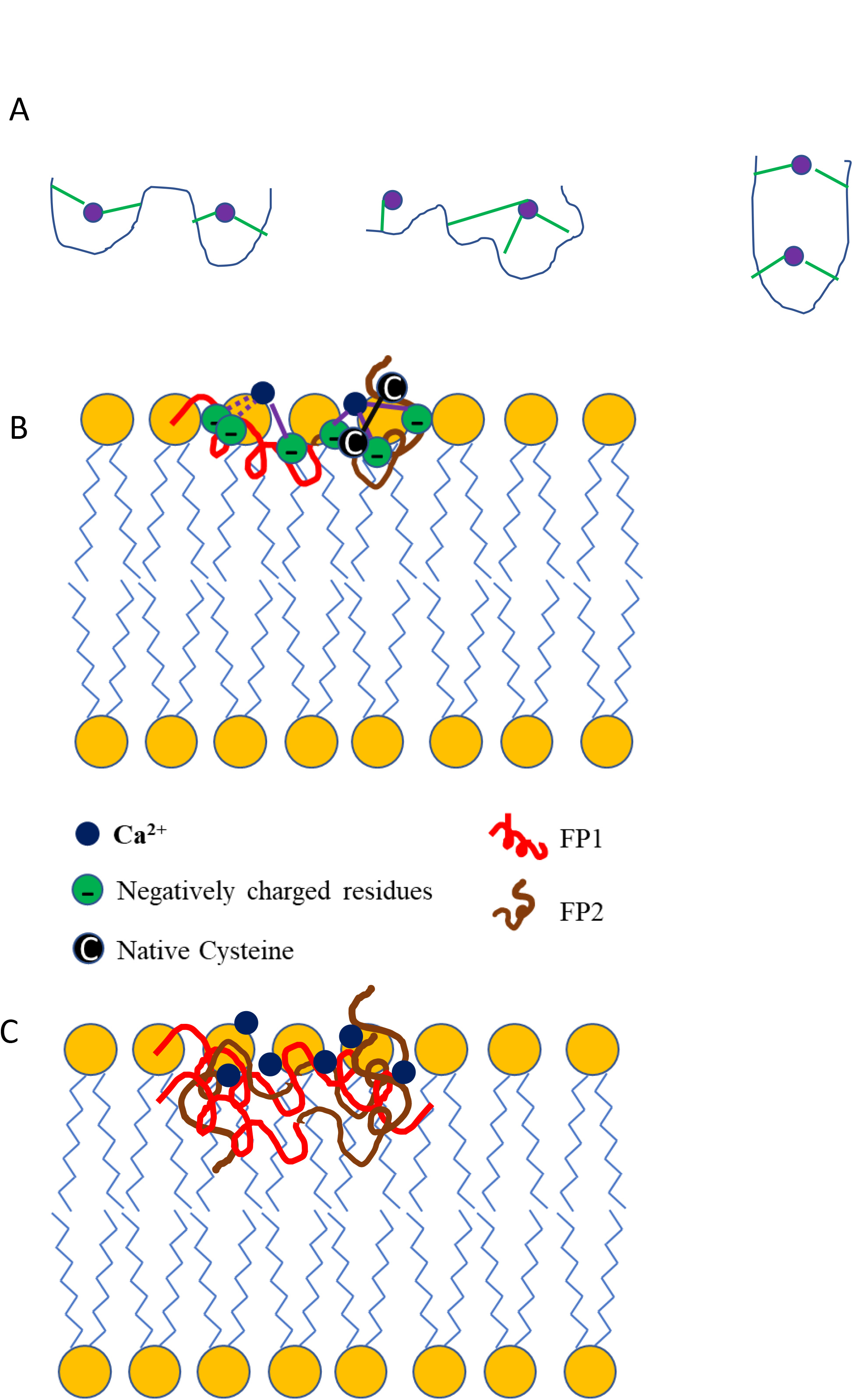
A) Different topologies of Ca^2+^ binding models. B) Model of the coronavirus FP interacting with a lipid bilayer. This model summarizes the data obtained in this work and highlights the unique features of this FP, including its bipartite nature and calcium-binding ability. C) Model of the trimerized FPs which inserts into the membrane deeper and has a greater effect on membrane ordering.

The reduction of ordering activity of the mutants investigated by ESR strongly favors the first topological model (Fig 10A, left): i.e., the three residues in the FP1 segment bind to one ion, and three residues in the FP2 segment bind to another. This model is further supported by the FP- Ca^2+^ binding titration (Figure 6A-D). We found that only the triple mutants in both FP1 and FP2 segments give a FP:Ca^2+^ binding ratio (N number) close to one. This is consistent with the first model, but contradicts the other models. If the second model (Fig 10B) is correct, we would expect the N number of E801A_D802A to be close to one: D812 binds the second ion but not the first ion, so this double mutant will lose its binding site for the first ion. However, its N number is 1.6. The same logic also excludes the possibility that E821 binds the first ion, as the N number of D825A_D831A is 1.58. If the third model is correct, then the triple mutants will still bind two ions, but their N number is close to one.

Of course, this FP-Ca^2+^ titration is in the solution, not in membranes. However, our ITC experiments on FP-membrane interaction support the first model from another aspect. The difference of partitioning enthalpy ΔH (Fig 7A and 7B), with or without Ca^2+^, indicates the folding of FP in the membrane due to the binding of Ca^2+^. Again, we find that the triple mutants have lowest ΔΔHs, indicating only one Ca^2+^ binding. And the three double mutants in the FP2 segments are very close to each other, indicating an even contribution in Ca^2+^ binding that is consistent with our ESR ordering activity results and FP-Ca^2+^ binding results. Comparing the FP- Ca^2+^ titration and the FP-membrane titration results, we found that although the former reaction occurs in solution and the latter in membranes, the FP-Ca^2+^ binding behaviors are largely consistent, i.e., the mutants with a lower N number in the FP-Ca^2+^ titration generally have a lower ΔΔH value in the FP-membrane titration (supplementary figure SF1). When the FP inserts into membranes, its overall accessibility to the Ca^2+^ should be different from that in the solution. However, SF1 shows that those negatively charged residues are fairly accessible to the Ca^2+^ even in membranes.

This model is somehow not consistent with a recent NMR study of the structure of SARS-2 FP^23^, which suggests that D830 (corresponding to D812 in SARS-1 FP) does not bind to Ca^2+^, and D839 and D843 (corresponding to E821 and D825 in SARS-1 FP) most likely participate in binding Ca^2+^, while the other three are moderated likely to bind Ca^2+^. This research is on SARS- 2 FP, so it may be a different from our SARS-1 FP case. More importantly, the environment used in the study consists of bicelles composed of mainly DMPC, which lacks the negatively charged headgroup such as PS used in our study. The negative charged headgroups attract Ca^2+^ to the membrane surface and thus increase the Ca^2+^ concentration in the headgroup region. Thus, the failure in attracting the Ca^2+^ by the neutral headgroup will significantly reduce the accessibility of the negatively charged residues of the FP to Ca^2+^, and thus lower the interaction between them. In that NMR study, the amide peaks of the backbone only shifted minimally, indicating that the interactions between the FP and Ca^2+^ are indeed minimal in that environment. The cholesterol used in our study also plays a role in the FP-Ca^2+^ interaction since cholesterol increases the fluidity of the membrane, and may affect the structure of FP in some fashion^48^. Although there has been no systematical study on the effect of cholesterol on the structure of SARS FPs, cholesterol is known to alter the structure of HIV FP significantly as well as having an effect on membranes.^10, 49^

### Why the FP-Ca^2+^ interaction is important for the membrane fusion?

Calcium ions are important modulators of membrane fusion. Because of their positive charges, they possess a generally enhancing or indirect effect on membrane fusion by electrostatic interactions with negatively charged headgroups of lipid bilayers. They thus decrease the electrostatic repulsion of two opposing membranes that are in close proximity prior to undergoing fusion. Calcium ions can also directly interact with fusion protein machineries, thereby activating their fusogenicity; and in such cases membrane fusion is clearly calcium-dependent. An example is the cellular SNARE-mediated synaptic vesicle fusion machinery^50–52^, i.e., without calcium ions present, membrane fusion does not take place. There are other situations in which calcium cations have been shown to interact with protein fusogens and enhance their fusogenicity; but without calcium, membrane fusion can still occur albeit at a reduced rate. In such cases membrane fusion is only partly dependent on calcium ions. The SARS-1, SARS-2^16^ and MERS FPs^15^ are in this category.

Ca^2+^ promotes the folding of SARS-1 FP in the membrane. First, from the CD spectra (Fig 4A), we can see an increase of helical content when the concentration of Ca^2+^ increases. Second, the FP-membrane titration results (Fig 7A and 7B) showing that in the presence of Ca^2+^, the FP folds better in the membrane. On the same time, Ca^2+^ also promotes the activity of FP in the Ca^2+^ response experiment (Fig 4B and 4C). Thus, the binding of Ca^2+^, the folding of FP in membranes, and the membrane ordering activity are strongly correlated.

In order to promote the folding of FP and the activity of FP, a bound Ca^2+^ ion needs to simultaneously bind to at least two binding sites. As we can see in the Ca^2+^-FP titration (Fig 6), even double mutants in either FP1 or FP2 have an N number close to 2, which means that even there is only one negatively charged residue remaining unsubstituted in either FP1 or FP2 segment, this residue can still bind one Ca^2+^. However, as shown by the ESR, all double mutants basically have a very small ordering activity at the same level of the triple mutants (Fig. 2C-D and Fig 3C-D). This indicates that at least two Ca^2+^ binding sites in each segment are required in order to maintain its “Ca^2+^-enhanced” ordering activity.

Thus, it seems that Ca^2+^ functions in two ways. The first way is to fix the FP in the interfacial region of the bilayer to help the formation of secondary structure of the FP in membrane. The second way is to promote the formation of tertiary structure of FP by holding two binding sites together. Both help the FP to posit in the membrane and squeeze the water molecules out of the headgroup region, and thus induce the ordering effect.

Previously, we found that the full-length SARS-1 FP (aka FP1_2) induces a greater membrane ordering effect than the sum of the combined individual FP1 and FP2. FP1_2’s ordering effect also reaches into the middle hydrophobic region (10PC), while neither FP1 nor FP2 has this ability. We have proposed that there is a synergetic effect because of the cooperation between the FP1 and FP2. Although the details of this cooperation are unknown, we suspected there is a FP1- FP2 interaction. As we have shown previously for the case of influenza HA, the interaction between FP-TMD increases its ability to perturb the membrane to a greater extent.^11^ Our research on HIV FPs also suggests that when HIV FPs are in the aggregation form, they induce a greater extent of membrane ordering as well as the effect reaches deeper^10^.

The recent NMR paper on SARS-2 FP structure mentioned above indicates that the FP inserts in the membrane as a wedge^23^, where the FP1 and FP2 are closely positioned together in membranes. Although that result is about the SARS-2 FP, it is very reasonable to believe that the SARS-1 FP can adopt a similar structure based on the similarity between the SARS-1 and SARS-2 FPs. And though we argue above that the Ca^2+^ binding topology in that research is different from ours because of the composition of lipids, it provides an insight into the possibility of interaction(s) between FP1 and FP2.

However, the limitation of the NMR and the MD simulation studies is that there is only one FP in their system: one FP in a bicelles in the NMR study^23^; or one FP in a membrane patch in the MD simulation^27^. These environments exclude the possibility of inter-molecular interaction between the FPs. However, this is possible in our ESR study system as the FPs bind to liposomal membranes which are large enough to harbor multiple FPs.

Considering in the “real biological scenario”, the three FPs are in a close position as they are in the whole S protein trimer. Thus, they are likely to form trimers as well. It is even possible that a higher number N-mer can be formed if multiple S protein trimers accumulate near a fusion site. It has been shown that in the case of influenza, 7-9 trimers accumulate in the fusion site^53, 54^. HIV FP is known to aggregate in the membrane and are supposed to aggregate in the context of whole gp41 timer during the membrane fusion as well^55^. However, we need a system that simulates the “real biological scenario” to test. That is why we develop a PP-SUV system.

### Does the separate FP work the same fashion as the FP on the whole S protein trimer?

It has long been questioned whether separate FPs function the same as in the whole protein. Our PP-SUV methodology monitors the membrane ordering induced by the pre-assembled S protein on the surface of the PP in real time. This provides several advantages. First, the FP is now a part of the entire protein. Second, the protein is pre-assembled, presumably in a trimer as suggested by Cryo-EM^17^. This allows the three FPs in the trimer to be located in close proximity, which is quite different than separate FP monomers. Third, the S protein trimers are now on a membrane of the enveloped particle, in which the spatial distribution is similar to that on the viral surface. This is a more “biological scenario” than the separate FP. In addition, by using time-resolved ESR we can monitor the change of S_0_ in real time, which allows us to study the kinetics of membrane ordering as well.

We have obtained some useful observations using this method. First, the FP on the PP induces membrane ordering of a magnitude equivalent to that of the separate FP (Fig 9E). This strongly suggests the “final” effect of FP on the PP and separate FP are similar.

Second, the mutations of the negatively charged residue affect both the extent and the kinetic of the ordering effect (Fig 9D) but the extent of the PP-induced ordering effect is similar to that of the corresponding separate FP. This confirms most results of our mutation study on the FP can be translated into the context of PP. The production of mutant S protein on PP is more difficult to control, as some mutants have very low expression levels or do not process well due to complicated reasons. Thus, the study on the FP mutants is useful.

Third, the activity of the whole protein is even more dependent on Ca^2+^ since the “basal activity” is minimal compared to that of the separate FPs (Fig 9C). But it is also possible that the “basal activity” will take longer time to take effect, which is out of our current acquisition time range. We need more experiment to test this hypothesis.

Fourth and most relevant to our previous discussion on the intermolecular interaction, in an initial experiment, we found that the individual FPs induce membrane ordering to a similar extent, but much more slowly (Fig 9F). Although careful control of relative quantities is required, it appears that the PP is more rapid in inducing the local ordering. This strongly suggests the existence of FP trimer and its benefit for the membrane fusion. The FPs are in close spatial location, since the S protein on the PP is pre-assembled as a trimer like the ones on the natural viral membranes, whereas the individual FPs are not. This suggests that initial trimerization of the FP helps to promote the local ordering. If this is the case, then an intermolecular FP1-FP2 interaction, i.e., the FP1 of one FP interacts with the FP2 of another FP is more likely to form. This will allow the FP1 and FP2 cooperate to induce stronger and deeper ordering effect as discussed above. It will require more research to confirm this hypothesis.

Although we are still improving our methodology, it has already shown potential for studying the initial stage of membrane fusion. In particular, we can monitor in real time how local membrane ordering develops in the SUV as mediated by the FPs in the complete glycoprotein trimers that are assembled on the PP surface. By comparing the kinetics of membrane ordering effects mediated by both the individual FP vs. the FP in the whole glycoprotein on PP membranes, we can expect to better understand this step in viral membrane fusion.

In addition to studying the mechanism, this system is also able to serve as a platform for testing the effects of environmental factors (pH, ions, etc.) and the inhibitory effects of drugs, peptides, and antibodies that target the FP. We show one example in Supplementary Figure SF2. We obtained a compound from our collaborator Dr Hector Aguilar-Carreno which has the potential to inhibit viral entry. We added the compound to the SARS-1 PP-SUV system. The compound does show a significant inhibitory effect on the membrane ordering, while the control version has much less inhibitory effect.

Although this present work mainly focuses on the SARS-1 FP, the knowledge is also useful for SARS-2 FP since their sequences are similar. We show the ESR results of two mutants as examples in Fig 8. Those mutants exhibit reduced ordering activity as do the corresponding mutants for SARS-1 FP. We also show initial results for the SARS-2 PP-SUV (Fig 9C), and it is clear that the system works for SARS-2 PP as well. Of course, as we have shown previously, although the SARS-2 and SARS-1 FPs differ only in three residues, the SARS-2 FP has a significant greater ordering effect^16^.

### The model of SARS FP in membranes

A proposed model for the SARS-1 FP, based on our observations and measurements, is shown in fig.10 B. The bipartite FP or “platform” consists of two distinct subdomains FP1 and FP2, each binding to a calcium ion via three negatively charged residues: aspartic acids (D) and glutamic acids (E). In the FP1 segment, D812 serves as a binding site, and the E801 and D802 serve together as another binding site and they are redundant; in the FP2 segment, E821, D825 and D831 serve as three binding sites. The loss of binding sites (i.e., by mutation) reduces its activity, especially when only one binding site is present. These Ca^2+^ bindings promote the folding of the FP in the membrane and enable the FP to insert more deeply into the shallower hydrophobic region (5PC). This insertion leads to the dehydration of the membrane and results in the membrane ordering. The possible trimerization of FPs (We draw this trimer model separately in fig. 10C, omitting the Ca^2+^ binding details as shown in fig 10B for simplicity) enables the cooperation of the FP1 and FP2 intermolecularly and thus enables the FP to induce the membrane ordering to a greater extent and deeper into the middle hydrophobic region (10PC)

## MATERIALS AND METHODS

### Lipids and Peptides

The lipids POPC, POPS, and the chain spin labels 5PC, 10PC and 14PC and a head group spin label dipalmitoylphospatidyl-tempo-choline (DPPTC) were purchased from Avanti (Alabaster, AL) or synthesized by our lab according to previous protocols. Cholesterol was purchased from Sigma (St. Louis, MO). All peptides were synthesized by BioMatik (Wilmington, DE) or SynBioSci Co. (Livermore, CA). The sequences of the peptides and the structure of the spin labeled lipids are shown in fig. 1D.

### Vesicle Preparation

The composition of membranes used in this study is consistent with our previous study ^41^. The desired amount of POPC, POPG, cholesterol and 0.5% (mol:mol) spin-labeled lipids in chloroform were mixed well and dried by N_2_ flow. The mixture was evacuated in a vacuum drier overnight to remove any trace of chloroform. To prepare MLVs, the lipids were resuspended and fully hydrated using 1 mL of pH 7 or pH 5 buffer (5 mM HEPES, 10 mM MES, 150 mM NaCl, and 0.1 mM EDTA, pH 7 or pH 5) at room temperature (RT) for 2 hours. To prepare SUVs for CD and ITC measurements and the PP-SUV system, the lipids were resuspended in pH 7 or pH 5 buffer and sonicated in an ice bath for 20 minutes or when the suspension became clear. The SUVs solution was then further clarified by ultracentrifugation at 13,000 rpm for 10 min.

### Circular Dichroism Spectroscopy

The CD experiments were carried out on an AVIV CD spectrometer Aviv Model 215. The peptides were mixed with SUVs in 1% P/L ratio with a final peptide concentration of 0.1 mg/mL at RT for more than 10 min before the measurements. The measurements were performed at 25 °C and two repetitions were collected. Blanks were subtracted and the resulting spectra were analyzed. The mean residue weight ellipticity was calculated using the formula: [Θ] = θ/(10×c×l×n), where θ is the ellipticity observed (in degrees); c is the peptide concentration (in dmol); l is the path length (0.1 cm); and n is the number of amino acids per peptide^45^. The percentages of secondary components were calculated using K2D3 online program^42^.

### Isothermal Titration Calorimetry

ITC experiments were performed in an N-ITC III calorimeter (TA Instrument, New Castle, DE). To measure the enthalpy of FP membrane binding, FP at 20 µM was injected into 1 mL 5 mM SUV solution at 37 °C. Each addition was 10 µL, each injection time was 15 sec, and each interval time was 5 min. Each experiment comprised about 25 to 30 injections. The data were analyzed with Origin (OriginLab Corp., Northampton, MA).

To measure the peptide-Ca^2+^ interaction, a total of 300 μL 2 mM CaCl_2_ in pH 5 buffer was injected into 0.4 mM FP in pH 5 buffer at 37 °C in a stepwise manner consisting of 10 µL per injection except that the first injection was 2 μL. The injection time was 15 sec for each injection and the interval time was 10 min. The background caused by the dilution of CaCl_2_ was subtracted by using data from a control experiment that titrated CaCl_2_ in a pH 5 buffer. The data were analyzed with Origin. The one-site model was used in the fitting to calculate the thermodynamic parameters. The protein concentration is determined by dry weight and UV spectroscopy at wavelength = 280 nm with extinction coefficient ε=1520 M^-1^ cm^-1^ for SARS-2 and SARS-1 FP in oxidized condition.

### ESR spectroscopy and nonlinear least-squares fit of ESR spectra

To prepare the samples for lipid ESR study, the desired amounts of FPs (1 mg/mL) were added into the lipid MLVs dispersion. After 20 min of incubation, the dispersion was spun at 13,000 rpm for 10 min. The concentrations of peptide were measured using UV to ensure complete binding of peptide. The pellet was transferred to a quartz capillary tube for ESR measurement. ESR spectra were collected on an ELEXSYS ESR spectrometer (Bruker Instruments, Billerica, MA) at X-band (9.5 GHz) at 25 °C using a N_2_ Temperature Controller (Bruker Instruments, Billerica, MA).

The ESR spectra from the labeled lipids were analyzed using the NLLS fitting program based on the stochastic Liouville equation ^29, 56^ using the MOMD or Microscopic Order Macroscopic Disorder model as in previous studies ^8–11, 14^. The fitting strategy is described below. We employed the Budil *et al.* NLLS fitting program ^29^ to obtain convergence to optimum parameters. The g-tensor and A-tensor parameters used in the simulations were determined from rigid limit spectra ^8^. In the simulation, we required a good fit with a small value of χ^2^ and also good agreement between the details of the final simulation and the experimental spectrum. Each experiment (and subsequent fit) was repeated 2 or 3 times to check reproducibility and estimate experimental uncertainty. Two sets of parameters that characterize the rotational diffusion of the nitroxide radical moiety in spin labels were generated. The first set is the rotational diffusion constants. R⊥ and R_||_ are respectively the rates of rotation of the nitroxide moiety around a molecular axis perpendicular and parallel to the preferential orienting axis of the acyl chain. The second set consists of the ordering tensor parameters, S_0_ and S_2_, which are defined as follows: S_0_=<D_2,00_>=<1/2(3cos2θ−1)>, and S_2_=<D_2,0_^2+^D_2,0-2_> =<√(3/2)sin2θcos2φ>, where D_2,00_, D_2,02_, and D_2,0-2_ are the Wigner rotation matrix elements and θ and φ are the polar and azimuthal angles for the orientation of the rotating axes of the nitroxide bonded to the lipid relative to the director of the bilayer, i.e., the preferential orientation of lipid molecules; the angular brackets imply ensemble averaging. S_0_ and its uncertainty were then calculated in standard fashion from its definition and the dimensionless ordering potentials C_20_ and C_22_ and their uncertainties found in the fitting. The typical uncertainties we find for S_0_ range from 1-5 × 10^-3^, while the uncertainties from repeated experiments are 5-8 × 10^-3^ or less than ±0.01. S_0_ indicates how strongly the chain segment to which the nitroxide is attached is aligned along the normal to the lipid bilayer, which is believed to be strongly correlated with hydration/dehydration of the lipid bilayers. As previously described, S_0_ is the main parameter for such studies ^11, 57, 58^.

### Pseudotyped virus particle – SUV docking experiments

The murine leukemia virus-based SARS-1-CoV S-pseudotyped particles, harboring wild-type (wt) SARS-CoV S protein, were generated using a replication deficient murine leukemia virus core in HEK-293T cell line as previously described ^59^ and provided by Gary Whittaker and Susan Daniel Labs. The Western blotting, cell viability assays and functional pseudotyped virus infectivity assays were also performed in Whittaker and Daniel Labs to ensure the expression and the activities of the S-proteins on the pp. The concentration of PP and the SUV with spin labeled lipids were determined by NanoSight NS300 (Malvern Panalytical, Malvern, UK). The PP were diluted to about 1× 10^6^ particle/ml and the SUV were mixed in roughly 1:5 ratio in the pH5 buffer at RT transferred in an ESR capillary. The mixture is triggered by the injection of concentrated CaCl_2_ to make the final Ca^2+^ concentration to 1 mM and immediately start the acquisition. The ESR spectra were collected at RT every 30 sec up to the duration of 15 min or longer. The ESR spectra were denoised using the wavelet denoising program5 as needed and the local S_0_ for each spectrum was extracted using the NLSL program^28^. The S_0_-time function were then plotted.

## Acknowledgments

This work was funded by NIH grants R01GM123779 and P41GM103521. We thank Tiffany Tang and Miya Kristine Bidon from the Dr. Susan Daniel Lab and the Dr. Gary Whittaker Lab for preparing the PP, for examining the expression level and measuring the concentrations. Dr. Daniel and Dr. Whittaker provided numerous helpful discussions and suggestions. We thank Dr. Brian Crane for the use of the CD spectrometer.

## Abbreviations

CD: circular dichroism spectroscopy
Chol: cholesterol
ITC: isothermal titration calorimetry
EPR: electron paramagnetic resonance spectroscopy
FP: fusion peptide
ITC: isothermal titration calorimetry
MERS: Middle East Respiratory Syndrome
MLV: multilamellar vesicle
MOMD: microscopic order macroscopic disorder
POPC: 1-palmitoyl-2-oleoyl-sn-glycero-3- phosphocholine
POPS: 1-palmitoyl-2-oleoyl-sn-glycero-3-phosphoserine
SARS: Severe Acute Respiratory Syndrome
SUV: small unilamellar vesicle
WT: wild type

---

**SF1.**
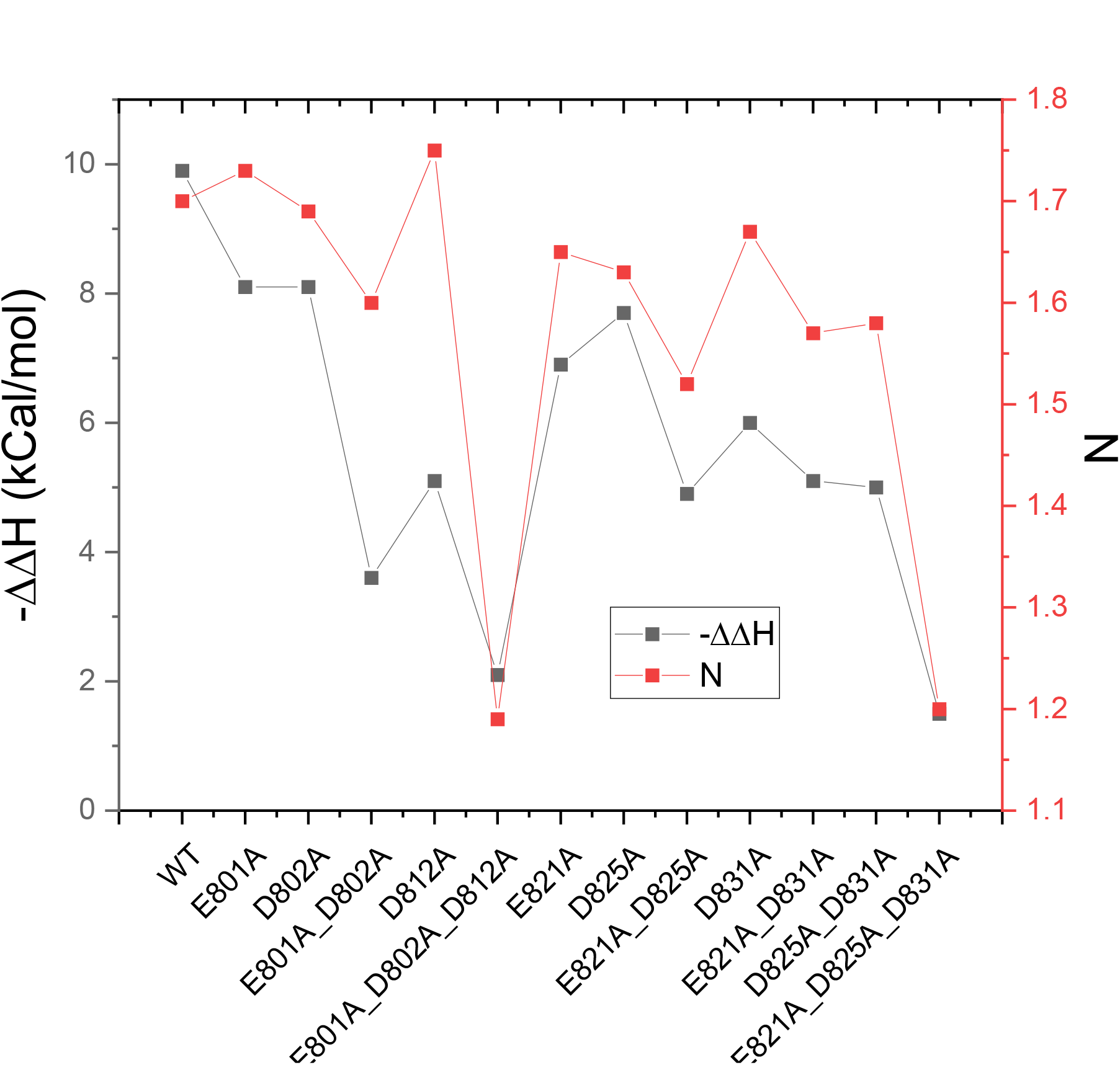
comparing the ΔΔH and N

**SF2.**
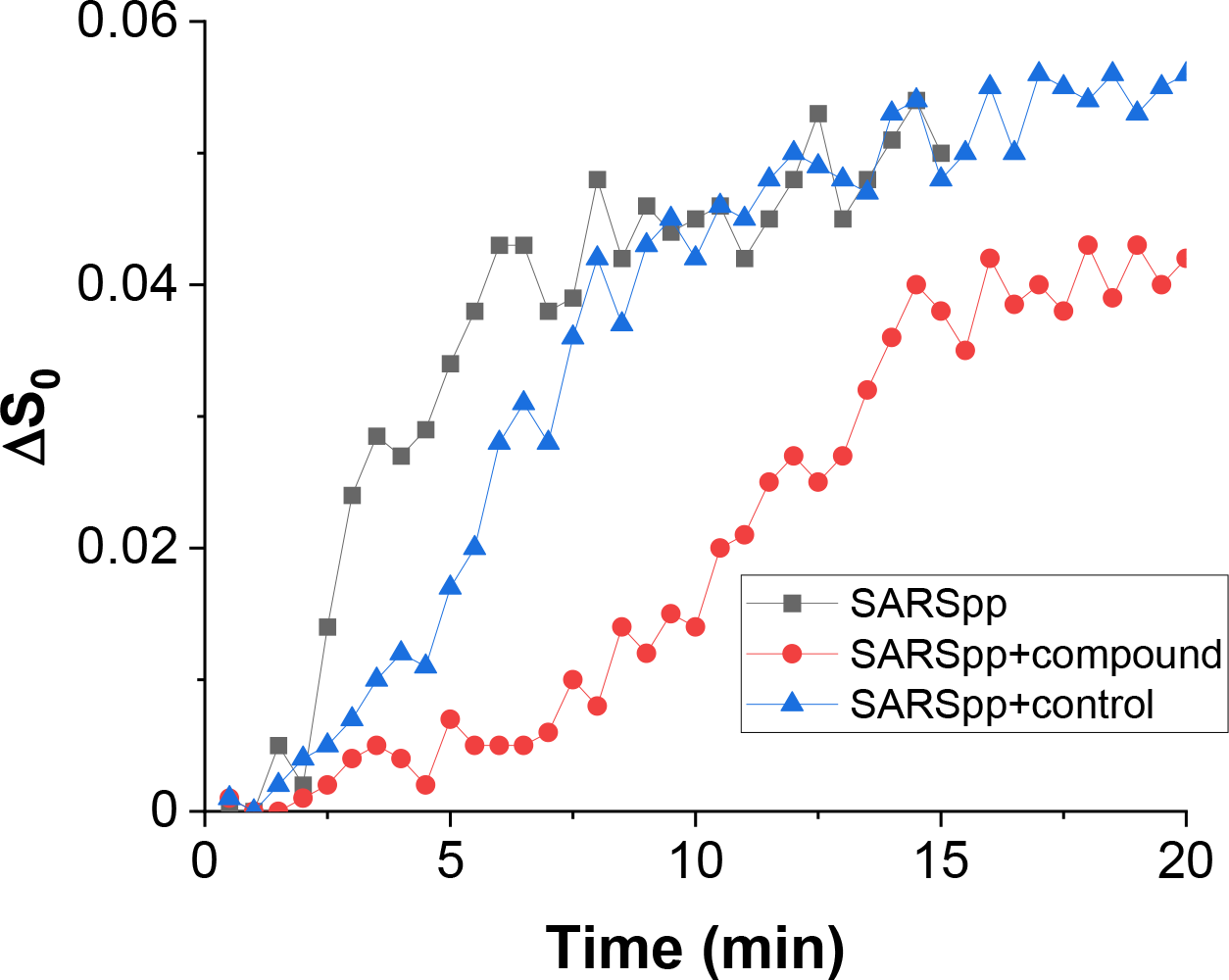
The effect of a compound on the pp mediated ordering in pp-SUV system

## Notes

### Competing Interest Statement

The authors have declared no competing interest.

## References

1. De Wit, E., Van Doremalen, N., Falzarano, D. & Munster, V. J. SARS and MERS: Recent insights into emerging coronaviruses. Nature Reviews Microbiology 14, 523–534 (2016).

2. Epand, R. M. Membrane fusion. Biosci Rep 20, 435–441 (2000).

3. Blumenthal, R., Clague, M. J., Durell, S. R. & Epand, R. M. Membrane fusion. Chem Rev 103, 53–69 (2003).

4. Weissenhorn, W. et al. Structural basis for membrane fusion by enveloped viruses. Mol. Membr. Biol. 16, 3–9 (1999).

5. Tang, T., Bidon, M., Jaimes, J. A., Whittaker, G. R. & Daniel, S. Coronavirus membrane fusion mechanism offers a potential target for antiviral development. Antiviral Res. 178, 104792 (2020).

6. White, J., Kartenbeck, J. & Helenius, A. Membrane fusion activity of influenza virus. Embo J 1, 217–222 (1982).

7. Sieczkarski, S. B. & Whittaker, G. R. Dissecting virus entry via endocytosis. J Gen Virol 83, 1535–1545 (2002).

8. Ge, M. & Freed, J. H. Fusion peptide from influenza hemagglutinin increases membrane surface order: an electron-spin resonance study. Biophys J 96, 4925–4934 (2009).

9. Ge, M. & Freed, J. H. Two conserved residues are important for inducing highly ordered membrane domains by the transmembrane domain of influenza hemagglutinin. Biophys J 100, 90–97 (2011).

10. Lai, A. L. & Freed, J. H. HIV gp41 fusion peptide increases membrane ordering in a cholesterol-dependent fashion. Biophys J 106, 172–181 (2014).

11. Lai, A. L. & Freed, J. H. Interaction between the influenza HA fusion peptide and transmembrane domain affects membrane structure. Biophys J. 109, 1–14 (2015).

12. Lai, A. L., Millet, J. K., Daniel, S., Freed, J. H. & Whittaker, G. R. The SARS-CoV Fusion Peptide Forms an Extended Bipartite Fusion Platform that Perturbs Membrane Order in a Calcium-Dependent Manner. J. Mol. Biol. 429, 3875–3892 (2017).

13. Nathan, L. et al. Calcium Ions Directly Interact with the Ebola Virus Fusion Peptide To Promote Structure–Function Changes That Enhance Infection. ACS Infect. Dis. 6, 250–260 (2020).

14. Pinello, J. F. et al. Structure-Function Studies Link Class II Viral Fusogens with the Ancestral Gamete Fusion Protein HAP2. Curr. Biol. 27, 651–660 (2017).

15. Straus, M. R. et al. Ca 2+ ions promote fusion of Middle East respiratory syndrome coronavirus with host cells and increase infectivity. J. Virol. 94, e00426–20 (2020).

16. Lai, A. L. & Freed, J. H. SARS-CoV-2 Fusion Peptide has a Greater Membrane Perturbating Effect than SARS-CoV with Highly Specific Dependence on Ca2+. J. Mol. Biol. (2021). doi:10.1016/j.jmb.2021.166946

17. Gui, M. et al. Cryo-electron microscopy structures of the SARS-CoV spike glycoprotein reveal a prerequisite conformational state for receptor binding. Cell Res. 27, 119–129 (2017).

18. Walls, A. C., et al. Structure, Function, and Antigenicity of the SARS-CoV-2 Spike Glycoprotein. Cell 181, 281–292.e6 (2020).

19. Wrapp, D. et al. Cryo-EM structure of the 2019-nCoV spike in the prefusion conformation. Science (80-.). (2020). doi:10.1126/science.aax0902

20. Cai, Y. et al. Distinct conformational states of SARS-CoV-2 spike protein. Science (80-.). (2020). doi:10.1126/science.abd4251

21. Xia, S. et al. Inhibition of SARS-CoV-2 (previously 2019-nCoV) infection by a highly potent pan-coronavirus fusion inhibitor targeting its spike protein that harbors a high capacity to mediate membrane fusion. Cell Res. 30, 343–355 (2020).

22. Wang, Q. et al. Structural and Functional Basis of SARS-CoV-2 Entry by Using Human ACE2. Cell (2020). doi:10.1016/j.cell.2020.03.045

23. Koppisetti, R. K., Fulcher, Y. G. & Van Doren, S. R. Fusion Peptide of SARS-CoV-2 Spike Rearranges into a Wedge Inserted in Bilayered Micelles. J. Am. Chem. Soc. (2021). doi:10.1021/jacs.1c05435

24. Georgieva, E. R., Ramlall, T. F., Borbat, P. P., Freed, J. H. & Eliezer, D. The lipid-binding domain of wild type and mutant α-synuclein: Compactness and interconversion between the broken and extended helix forms. J. Biol. Chem. 285, 28261–28274 (2010).

25. Georgieva, E. R., Xiao, S., Borbat, P. P., Freed, J. H. & Eliezer, D. Tau binds to lipid membrane surfaces via short amphipathic helices located in its microtubule-binding repeats. Biophys. J. 107, 1441–1452 (2014).

26. Snead, D. et al. Unique structural features of membrane-bound C-terminal domain motifs modulate complexin inhibitory function. Front. Mol. Neurosci. 10, 1–17 (2017).

27. Khelashvili, G., Plante, A., Doktorova, M. & Weinstein, H. Ca2+-dependent mechanism of membrane insertion and destabilization by the SARS-CoV-2 fusion peptide. Biophys J. in press, 2020.12.03.410472 (2021).

28. Ge, M., Budil, D. E. & Freed, J. H. An electron spin resonance study of interactions between phosphatidylcholine and phosphatidylserine in oriented membranes. Biophys J 66, 1515–1521 (1994).

29. Budil, D. E., Lee, S., Saxena, S. & Freed, J. H. Nonlinear-least-squares analysis of slow-motion EPR spectra in one and two dimensions using a modified Levenberg-Marquardt algorithm. J. Magn. Reson. Ser. A 120, 155–189 (1996).

30. Lou, Y., Ge, M. & Freed, J. A Multifrequency ESR Study of the Complex Dynamics of Membranes. J Phys Chem B 105, 11053–11056 (2001).

31. Ge, M. & Freed, J. H. Hydration, structure, and molecular interactions in the headgroup region of dioleoylphosphatidylcholine bilayers: an electron spin resonance study. Biophys J 85, 4023–4040 (2003).

32. van Meer, G., Voelker, D. R. & Feigenson, G. W. Membrane lipids: where they are and how they behave. Nat. Rev. Mol. Cell Biol. 9, 112–124 (2008).

33. Lai, A. L., Tamm, L. K., Ellena, J. F. & Cafiso, D. S. Synaptotagmin 1 modulates lipid acyl chain order in lipid bilayers by demixing phosphatidylserine. J Biol Chem 286, 25291–25300 (2011).

34. Takamori, S. et al. Molecular anatomy of a trafficking organelle. Cell 127, 831–846 (2006).

35. Frazier, A. A., Roller, C. R., Havelka, J. J., Hinderliter, A. & Cafiso, D. S. Membrane-bound orientation and position of the synaptotagmin I C2A domain by site-directed spin labeling. Biochemistry 42, 96–105 (2003).

36. Patel, A., Mohl, B.-P. & Roy, P. Entry of Bluetongue Virus Capsid Requires the Late Endosome-specific Lipid Lysobisphosphatidic Acid. J. Biol. Chem. 291, 12408–19 (2016).

37. Matos, P. M. et al. Anionic lipids are required for vesicular stomatitis virus G protein-mediated single particle fusion with supported lipid bilayers. J. Biol. Chem. 288, 12416– 25 (2013).

38. Zaitseva, E. et al. Fusion Stage of HIV-1 Entry Depends on Virus-Induced Cell Surface Exposure of Phosphatidylserine. Cell Host Microbe 22, 99–110.e7 (2017).

39. Hu, Y. & Patel, S. Thermodynamics of cell-penetrating HIV1 TAT peptide insertion into PC/PS/CHOL model bilayers through transmembrane pores: the roles of cholesterol and anionic lipids. Soft Matter 12, 6716–27 (2016).

40. Yang, R., Prorok, M., Castellino, F. J. & Weliky, D. P. A trimeric HIV-1 fusion peptide construct which does not self-associate in aqueous solution and which has 15-fold higher membrane fusion rate. J Am Chem Soc 126, 14722–14723 (2004).

41. Madu, I. G., Roth, S. L., Belouzard, S. & Whittaker, G. R. Characterization of a highly conserved domain within the severe acute respiratory syndrome coronavirus spike protein S2 domain with characteristics of a viral fusion peptide. J Virol 83, 7411–7421 (2009).

42. Louis-Jeune, C., Andrade-Navarro, M. A. & Perez-Iratxeta, C. Prediction of protein secondary structure from circular dichroism using theoretically derived spectra. Proteins 80, 374–381 (2012).

43. Brennan, S. C. et al. The extracellular calcium-sensing receptor regulates human fetal lung development via CFTR. Sci. Rep. 6, 21975 (2016).

44. Seelig, J., Nebel, S., Ganz, P. & Bruns, C. Electrostatic and Nonpolar Peptide-Membrane Interactions - Lipid-Binding and Functional-Properties of Somatostatin Analogs of Charge Z = +1 to Z = +3. Biochemistry 32, 9714–9721 (1993).

45. Lai, A. L., Park, H., White, J. M. & Tamm, L. K. Fusion peptide of influenza hemagglutinin requires a fixed angle boomerang structure for activity. J Biol Chem 281, 5760–5770 (2006).

46. Li, Y., Han, X. & Tamm, L. K. Thermodynamics of fusion peptide-membrane interactions. Biochemistry 42, 7245–7251 (2003).

47. Lai, A. L. & Freed, J. H. Influenza Fusion Peptide and Transmembrane Domain Interaction Induces Distinct Domains in Lipid Bilayers. Biophys. J. 106, 707a (2014).

48. Zakany, F., Kovacs, T., Panyi, G. & Varga, Z. Direct and indirect cholesterol effects on membrane proteins with special focus on potassium channels. Biochim. Biophys. Acta - Mol. Cell Biol. Lipids 1865, 158706 (2020).

49. Lai, A. L., Moorthy, A. E., Li, Y. & Tamm, L. K. Fusion activity of HIV gp41 fusion domain is related to its secondary structure and depth of membrane insertion in a cholesterol-dependent fashion. J Mol Biol 418, 3–15 (2012).

50. Bhalla, A., Chicka, M. C., Tucker, W. C. & Chapman, E. R. Ca(2+)-synaptotagmin directly regulates t-SNARE function during reconstituted membrane fusion. Nat Struct Mol Biol 13, 323–330 (2006).

51. Brunger, A. T. Structure and function of SNARE and SNARE-interacting proteins. Q Rev Biophys 38, 1–47 (2005).

52. Chicka, M. C., Hui, E., Liu, H. & Chapman, E. R. Synaptotagmin arrests the SNARE complex before triggering fast, efficient membrane fusion in response to Ca2+. Nat Struct Mol Biol 15, 827–835 (2008).

53. Podbilewicz, B. Virus and cell fusion mechanisms. Annu Rev Cell Dev Biol 30, 111–139 (2014).

54. Bentz, J. & Mittal, A. Architecture of the influenza hemagglutinin membrane fusion site. Biochim Biophys Acta 1614, 24–35 (2003).

55. Qiang, W., Bodner, M. L. & Weliky, D. P. Solid-state NMR spectroscopy of human immunodeficiency virus fusion peptides associated with host-cell-like membranes: 2D correlation spectra and distance measurements support a fully extended conformation and models for specific antiparallel strand regis. J Am Chem Soc 130, 5459–5471 (2008).

56. Ge, M., Budil, D. E. & Freed, J. H. ESR studies of spin-labeled membranes aligned by isopotential spin-dry ultracentrifugation: lipid-protein interactions. Biophys J 67, 2326–2344 (1994).

57. Ge, M. et al. Electron spin resonance characterization of liquid ordered phase of detergent-resistant membranes from RBL-2H3 cells. Biophys J 77, 925–933 (1999).

58. Ge, M. et al. Ordered and disordered phases coexist in plasma membrane vesicles of RBL-2H3 mast cells. An ESR study. Biophys J 85, 1278–1288 (2003).

59. Millet, J. K. & Whittaker, G. R. Murine Leukemia Virus (MLV)-based Coronavirus Spike-pseudotyped Particle Production and Infection. Bio-protocol 6, (2016).

